# Single-cell multi-omic analyses resolve the cellular diversity of ALK/ROS1/MET/NTRK-fused gliomas in infants and older children

**DOI:** 10.64898/2026.07.16.738862

**Authors:** Andrea J. De Micheli, Carlos O.A. de Biagi-Junior, Costanza Lo Cascio, Charbel Machaalani, Andreas Postlmayr, Shashank Katiyar, Roeltje R. Maas, Venkatesh Kancherla, Regina Reimann, Michal Zápotocký, Liana F. Nobre, Sebastian K. Eder, Jacob S. Rozowsky, Théo Ribierre, Fides Zenk, Adam Resnick, Matthew Clarke, Johannes Gojo, Uri Tabori, Cynthia Hawkins, Chris Jones, Florence M. G. Cavalli, Mariella G. Filbin, Ana S. Guerreiro Stücklin

**Affiliations:** Division of Oncology and Children’s Research Center, University Children’s Hospital of Zurich, Zurich, Switzerland; Department of Pediatric Oncology, Dana-Farber Boston Children’s Cancer and Blood Disorders Center, Boston, Massachusetts, USA; Institute for Neuropathology, University Hospital Zurich, Zurich, Switzerland; Department of Pediatric Hematology and Oncology, Second Faculty of Medicine, Charles University and University Hospital Motol, Prague, Czech Republic; Department of Paediatrics, University of Alberta, Edmonton, Alberta, Canada; Neuropathology Department, Great Ormond Street Hospital for Children NHS Foundation Trust, United Kingdom; NeuroNA Human Cellular Neuroscience Platform, Campus Biotech, Geneva, Switzerland; Department of Basic Neurosciences, University of Geneva, Geneva, Switzerland; Epigenomics of Neurodevelopment, Brain Mind Institute, School of Life Sciences, Ecole Polytechnique Federale de Lausanne, Ecublens, Switzerland; Center for Data-Driven Discovery in Biomedicine, Children’s Hospital of Philadelphia, Philadelphia, PA, United States; Division of Neurosurgery, Children’s Hospital of Philadelphia, Philadelphia, PA, United States; Division of Cancer Biology, Institute of Cancer Research, United Kingdom; Department of Pediatrics and Adolescent Medicine, Comprehensive Cancer Center and Comprehensive Center for Pediatrics, Medical University of Vienna, Vienna, Austria; St. Anna Children’s Hospital, Department of Pediatrics and Adolescent Medicine, Medical University of Vienna and St. Anna Children’s Cancer Research Institute (CCRI), 1090 Vienna, Austria; Arthur and Sonia Labbatt Brain Tumour Research Centre, The Hospital for Sick Children, Toronto, Ontario, Canada; Division of Molecular Pathology, The Institute of Cancer Research, London, UK; Institut National de la Santé et de la Recherche Médicale (INSERM), U1331 Computational Oncology, Paris, France; Institut Curie, PSL University, Paris, France; Mines Paris, PSL University, CBIO-Centre for Computational Biology, Paris, France

## Abstract

Pediatric cancers are thought to arise from dysregulation of developmental programs, otherwise tightly regulated in time and space. Infant-type hemispheric gliomas (IHGs) arise in early childhood, driven by characteristic ALK/ROS1/MET/NTRK receptor tyrosine kinase (RTK) gene fusions. We dissected the cellular hierarchies of 24 fusion-positive gliomas, spanning infants through adolescents, using single-cell and single-nucleus RNA/ATAC-seq, and spatial transcriptomics. We identified five cancer cell states, with radial glia-like cells at the apex of a neoplastic hierarchy resembling neuronal- and glial-like trajectories. Neuronal-like cells were enriched in most IHGs but diminished in ROS1-fused IHGs and older patients. Integration of chromatin profiling revealed FOS/JUN-driven oncogenic programs and high inferred plasticity across all cancer cell populations. Myeloid cells, the most abundant non-neoplastic population, comprised distinct subgroups, suggesting context-dependent functions. Despite lacking high-order structure, spatial transcriptomics revealed discrete cellular niches within IHGs. Collectively, our findings elucidate the cellular states and developmental programs underlying IHGs and RTK-fused gliomas in older patients, opening new avenues for research and therapy innovation.

## INTRODUCTION

Pediatric brain tumors are thought to stem from oncogenic events that disrupt cell maturation programs during neural development. Gliomas, the most common primary brain tumors across all age groups^1–5^, manifest in a molecular-, age- and location-defined fashion^3,4,6^. Pediatric high-grade gliomas (pHGGs), for example, arise in the brain hemispheres in infants and are driven by oncogenic fusions in receptor tyrosine kinase (RTK) genes. In contrast, in school-aged children, H3K27-altered diffuse midline gliomas (DMG) are more prevalent^7– 10^, whereas in adolescents and young adults, H3G34-mutated hemispheric tumors begin to emerge^11,12^. Molecular alterations characteristic of adult-type high-grade gliomas, including *IDH*^13^, *TERT* promoter^14,15^, and *EGFR* mutations, are uncommon in pediatric gliomas^16^.

In neonates and young children, a rare glioma subtype known as *infant-type hemispheric glioma* (IHG) is driven by oncogenic fusions involving the RTK genes *ALK, ROS1, MET*, or *NTRK1/2/3*^3,4,17^. These RTK fusions are characteristic but not exclusive to IHGs and can be detected across clinically diverse pediatric gliomas (henceforth RTK-fused gliomas), spanning low- to high-grade morphologies, suggesting heterogeneity in their cellular composition. In the most recent 5^th^ edition of the WHO classification of central nervous system (CNS) tumors^17^, IHGs are included as a distinct group of pediatric-type high-grade gliomas. However, they display low- to high-grade features and carry a more favorable prognosis than other pHGG^3,4^.

The distinctive molecular features of pediatric gliomas illustrate a broader paradigm tying their origins to disrupted developmental processes^8,18–20^. For instance, H3K27M-altered DMGs harbor cancer stem-like cells resembling oligodendrocyte progenitor cells (OPCs), whereas pediatric-type diffuse high-grade gliomas with H3G34R/V mutations arise from radial glia (RG)-like progenitors and follow an interneuron-like developmental trajectory^7,11^. In this context, key questions concerning the biology of IHGs and other RTK-fused gliomas remain unanswered: which cell populations and developmental programs are retained within these tumors? How do different RTK fusions impact developmental trajectories? How do these changes vary across age groups? What is the spatial cellular architecture of IHGs?

To address these questions, we profiled a cohort of 24 pediatric gliomas, selected for the presence of oncogenic RTK fusions (IHGs and older patients). Using single-cell and -nucleus RNA-sequencing as well as single-nucleus ATAC-sequencing, we resolved manifold cancer cell states retaining features of normal neural development. Our analyses reveal that IHGs harbor tumor cell states within a hierarchy rooted in RG-like progenitors, with further cancer cell populations resembling both neuronal- and glial-like progenitor trajectories. All tumor cell populations retain expression of early progenitor markers, suggesting a maturation block. Intermediate neuronal progenitor (iNPC)/Neuronal-like cells are enriched in most IHGs but reduced or absent in ROS1-driven IHGs and other RTK fusion-positive gliomas in older patients, suggesting a possible age-, fusion-, and/or lineage-dependent window of vulnerability that impacts tumor cell composition. Integration of single-nucleus RNA-seq and ATAC-seq data revealed high cellular plasticity and enrichment of FOS/JUN transcription factor activity, regulated downstream of MAPK signaling, in all cancer cell populations. Finally, spatial mapping revealed a lack of large-scale structural organization but identified localized niches where iNPC/Neuronal-like and glial progenitor cell/astrocyte (GPC/AC)-like populations form discrete spatial clusters. Together, our multi-omic analyses define the cellular hierarchies and spatial architecture of IHGs and other RTK-fused pediatric gliomas, providing new insights into tumor biology with clinical implications.

## RESULTS

### Cohort of IHG and other RTK-fused gliomas

We selected a cohort of 24 pediatric gliomas harboring a fusion in either one of the RTK genes of interest: *ALK* (n=7), *ROS1* (n=5), *NTRK* family (n=7), and *MET* (n=5). Samples were obtained from patients diagnosed between 0.23 and 15.5 years of age (median 3.16), with 14 patients diagnosed below the age of 3 (Infants), 6 between the ages of 3 and 13 years (Children), and 4 above the age of 13 (Adolescents) (Fig. 1A). Twenty-three samples originated from tumors located in the hemispheric regions and 1 sample from a tumor located in the cerebellum. Based on histopathological annotation at the time of initial diagnosis, 17 tumors were classified as high-grade gliomas (HGGs) and 7 as low-grade gliomas (LGGs). Methylation-based classification information was available for 16 patients, including 9/14 patients under 3 years of age. Within this infant group, methylation classification confirmed the diagnosis of IHG in 7/9 cases for which methylation was available (median age 1.0), one *EML4::ALK*-positive tumor in a 1.26-year-old patient classified as a low-grade glial/glioneuronal/neuroepithelial tumor, and one tumor was non-classifiable. For 5/14 infant tumors methylation data were not available but features were consistent with the diagnosis of IHG as per the most recent WHO classification of CNS tumors^17^ (Table S1).

**Figure 1.**
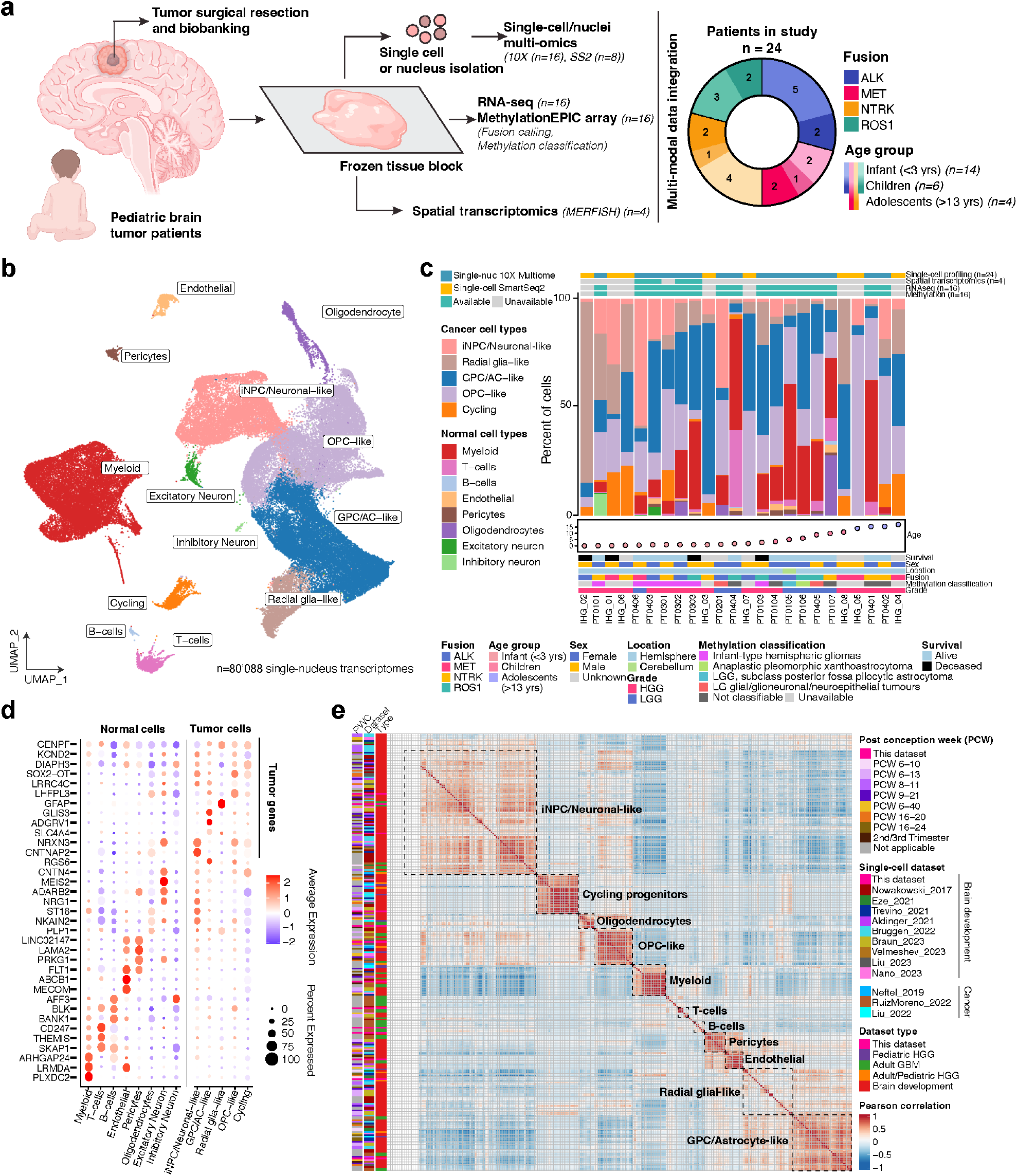
Cohort and single-nucleus atlas of IHGs and other RTK-fused pediatric gliomas. (a) Overview of study design: Frozen tumor biopsies were obtained from surgical resections and profiled by multi-omic analyses. Twenty-four patients were included in the study, grouped by fusion genes (ALK/ROS1/NTRKs/MET) and age (infants, children, adolescents). (b) Integrated UMAP of all single-nucleus transcriptomes classified into 8 normal and 5 cancer cell populations. (c) Proportion of non-tumor and tumor cell populations resolved by single-nucleus RNA-seq, and proportion of cancer cell population resolved by Single-cell SmartSeq2 organized by age. Upper legend indicates omics profiling method (MERFISH: n=4, 10X Multiome: n=16, SmartSeq2: n=8, RNA-seq: n=18, Methylation: n=16). Lower legend indicates fusion type, age group, sex, anatomical location, methylation class, tumor grade, and survival status. (d) Dot plot showing the average normalized expression and percent expressed in cluster for top differentially expressed genes. Cell populations and genes are divided into normal and cancer cell groups. (e) Heatmap showing metaprogram Pearson correlation analysis integrating gene signatures from the resolved populations (in b) with gene signatures from published single-cell datasets of the developing brain and adult glioma.

We profiled the cohort using single-nucleus, single-cell, and bulk RNA-sequencing methods (Fig. 1A). From frozen tissue, we isolated nuclei from 16 samples for joint single-nucleus RNA-seq and ATAC-seq (10X Multiome, Fig. S1A-E), and for the other 8 cases, we performed full-length single-cell RNA-seq (Smart-seq2^21^). Bulk RNA-seq identified fusion genes and their partners. Most fusions were unique, with only *PPP1CB::ALK*^22^ and *PTPRZ1::MET*^23^ recurring in two and three patients, respectively (Fig. S1F, Table S1). Finally, for 4/7 IHGs in this cohort, tumor sections were used for spatial transcriptomics (MERFISH)^24^, and 4 additional IHG tumor sections were profiled using Xenium. Collectively, we profiled IHG and other RTK-fused gliomas spanning diverse ages, RTK fusions, and tumor subtypes.

### Single-nucleus RNA-seq atlas of RTK-fused gliomas

To resolve the cellular diversity of RTK-fused gliomas, we assembled a snRNA-seq atlas (10X Multiome) and measured simultaneously the transcriptomes and chromatin accessibility of 80’088 cells from 16 patients, including 9 infants, of which 7 were methylation-confirmed IHGs (Fig. 1B, C and S1A-G). The unintegrated UMAP revealed distinct, patient-specific transcriptomic profiles and significant inter-patient heterogeneity among cancer cells, unlike most non-neoplastic cells (immune and vascular), which clustered together (Fig. S1C-E).

To reveal conserved gene expression programs, we generated an integrated atlas by using reciprocal PCA (RPCA) to correct for patient-specific signatures, based on the top 5000 most variable genes shared between patients (Fig. 1B, Fig. S1D). Unsupervised clustering identified a total of 13 groups of cells, with cell types typically found in pediatric and adult gliomas^7,8,11,25,26^, including 8 non-neoplastic and 5 cancer cell populations. We defined non-neoplastic cells based on each population’s top differentially expressed genes, which aligned with known cell markers, such as *LAMA2*+ pericytes (0.8%), *NRG1*+ excitatory and inhibitory neurons (1.3%), *FLT1*+ (*VEGFR1*+) endothelial cells (1.3%), *PLP1*+ oligodendrocytes (2.4%), *AFF3, BLK, BANK1* B-cells (0.2%), *CD247, THEMIS, SKAP1* T-cells (2%), *and ARHGAP24, LRMDA, PLXDC2* myeloid cells (22%) (Fig. 1C-D, Table S2).

Copy number variation (CNV) inference analyses further supported the identification of cancer cell states (Fig. S2). Next, we annotated each cancer cell population by correlating its differential gene expression profile to 265 distinct signatures drawn from 9 human fetal brain atlases^27–35^ and 3 pediatric/adult glioma datasets ^7,25,36^ (Fig. S1H, Table S3). This analysis identified cancer cell populations within discrete cell-type-specific gene expression modules (Fig. 1E, Fig. S3A-B), covering the cell states found in the developing brain, as well as immune and vascular cells (Fig. S3A-B). Through consensus across multiple reference correlations, we annotated five cancer cell populations as: *GFAP+ CLU+* Radial glia (RG)-like, *ADGRV1*+ *GLIS3*+ Glial progenitor cell (GPC)/Astrocyte (AC)-like, *CNTNAP2+ NRXN3+* intermediate Neuronal progenitor cell (iNPC)/Neuronal-like, *LHFPL3+* OPC-like, and *CENPF+* Cycling cells (Fig. 1D-E). Of note, we did not detect a clear mesenchymal-like (MES-like) population, as described in other pediatric and adult gliomas ^26,36,37^ . Metaprogram integration with existing datasets revealed a weak-to-moderate correlation between the MES-like and the RG-like and GPC/AC-like signatures, suggesting that different cells can co-opt mesenchymal programs (Fig S4).

### Cancer cell states retain features of normal neural development

We next further characterized the cancer cell states (n=69’305 cells, Fig. 2A). First, we compared their gene expression profiles to those of cells detected during human telencephalic development (post-conception weeks 5.85 to 37)^27^ (Fig. 2B). Analyses of expression scores revealed that cancer cell populations preferentially map to cell lineages of the developing brain: RG-like cells mapped mostly to early radial glia [RG-early], GPC/AC-like mapped to more differentiated ventricular, truncated, and outer radial glia cells [vRG, tRG, oRG] as well as astrocytes (Fig 2B, Fig. S5C). iNPC/Neuronal-like cells mapped to a broad range of intermediate neuronal progenitor cells from newborn excitatory neurons [IPC-nEN1-3], interneurons from medial and caudal ganglionic eminence [IN-CTX-MGE/CGE1-2], [nEN-early], interneurons [nIN1-5], and excitatory as well as newborn neurons [nEN-early], interneurons [nIN1-5], and excitatory neurons [EN]. Cycling progenitors mapped to radial glia and to cycling intermediate precursors, whereas OPC-like cells mapped with normal OPCs (Fig. 2B). Interestingly, RG-like cells also showed high expression scores for mature astrocytes and neuronal lineage cells. Mapping the expression of key brain development markers (Fig. 2C-D), we detected stem cell markers SOX2 and NES in all cancer cell states, with higher expression in RG-like cells. Radial glia markers (FABP7, GFAP) were most enriched in RG-like cells and expressed to different extents in all other populations. Similarly, OPC marker (OLIG2, SOX10, PDGFRA) expression concentrated in OPC-like cells and the astrocytic marker AQP4 in RG-like and GPC/AC-like populations. Immature and mature neuronal markers (DCX, RBFOX3/NeuN, SYN1, KCNJ6) were enriched in iNPC/Neuronal-like cells. While neural development lineage markers are expressed in cancer cells, their expression patterns are less defined than in normal brain tissue, suggesting inherent plasticity and lack of lineage commitment. Interestingly, a subset of RG-like cells expressed markers from all developmental lineages, suggesting a stem-like phenotype primed for differentiation (Fig. 2B).

**Figure 2.**
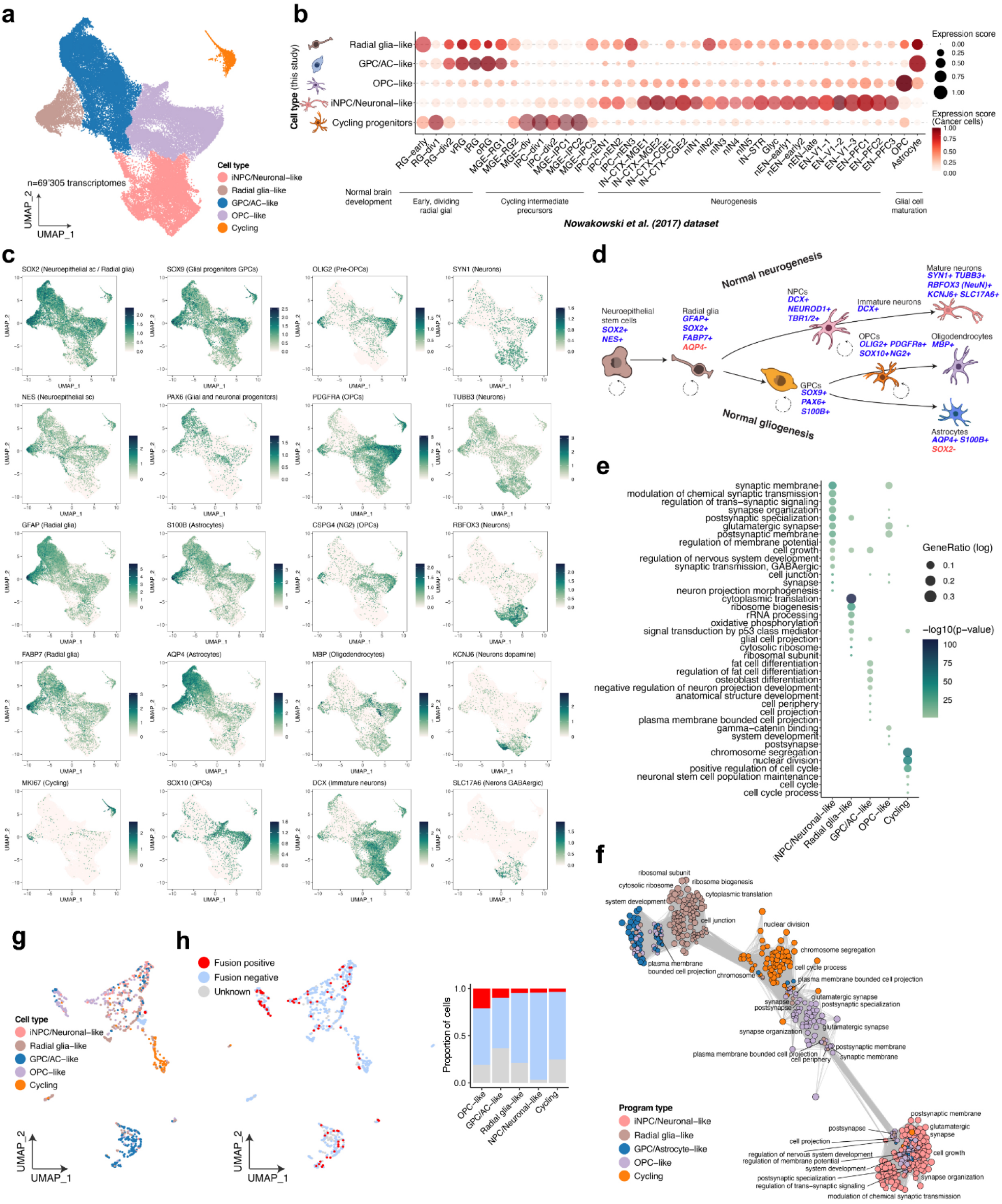
SnRNA-seq analyses reveal 5 cancer cell populations with developmental signatures. (a) Integrated UMAP of cancer cell populations resolved by snRNA-seq. (b) Projection of cancer cell gene expression signatures onto a single-cell atlas27 of the developing human cortex. Color scale: scores of normal cell signatures in cancer cells. Dot sizes: scores of cancer cell signatures in normal cells. (c) Expression patterns of key brain development marker genes. (d) Simplified representation of normal neurogenesis and gliogenesis pathways with selected key markers. (e) Dot plot of gene ontology (GO) terms, showing significance (-log10(p-value)) and gene ratio (proportion of genes associated with each term) across different cancer cell populations. (f) GO term enrichment network for genes upregulated in each of the 5 cancer cell populations and colored accordingly. Nodes represent enriched GO terms, with similar terms clustering together. The size of each node reflects the number of enriched genes for each term. (g) NMF-integrated UMAP of 877 cancer single-cell transcriptomes profiled by SmartSeq2, colored by cancer cell types. (h) UMAP of SmartSeq2 cells with gene fusion information. Fusion-positive corresponds to cells in fusion-positive samples with at least one fused transcript detected. Fusion negative corresponds to cells in fusion-positive samples for which no fused transcripts are detected. Unknown corresponds to cells for which no fused transcripts were detected in SmartSeq2 data, but the fusion was detected with another clinical assay.

Gene ontology analysis confirmed functional heterogeneity of these cancer cell states (Fig. 2E,F). Enrichment for terms related to translation, cell growth, and ribosome biogenesis in RG-like cells supports their stem-like self-renewing capabilities^38^ . In contrast, GPC/AC-like cells were enriched for glial differentiation terms, while both OPC-like and iNPC/Neuronal-like cells showed enrichment for synaptic specialization and signaling terms, such as *glutamatergic synapse*, which were identified in both iNPC/Neuronal-like and OPC-like cells, similar to previous reports^39^ (Fig. 2E, F).

We also used SmartSeq2 to generate an annotated full-length scRNA-seq dataset from 8 patients (n=877 cells) (Figs. 2G,H and S5A,B) and directly identified corresponding fusion transcripts within single cells (Fig. 2H). Fusion-positive reads were detected in 5/8 cases, confirming the initial diagnosis. Importantly, fusion transcripts were identified within all five cancer cell populations, including in RG-like cells, without enrichment of the fusion in a specific cell type (Fig. 2H).

In summary, RTK-fused gliomas comprise five transcriptionally distinct cancer cell states that retain key markers and functional programs of normal brain development but lack lineage commitment, with diffuse expression of early neural progenitor markers and fusion transcripts across all cancer cell populations.

### Developmental trajectory analysis of the glioma cancer cell compartment

To investigate whether RTK-fused glioma cells follow developmental trajectories resembling those of normal brain development, we used RNA velocity^40^ as a hypothesis-generating framework to explore transcriptional trends across cancer cell states (Fig. 3A). RG-like and a subset of GPC/AC-like cells occupied the top of the inferred hierarchy, with possible bidirectional transitions between them. Of note, these populations at the apex of the tumor hierarchy (RG-like and GPC/AC-like) share greater transcriptomic similarity with each other than with the other malignant cell states. From these populations, a putative trajectory suggested two broad branches: OPC-like and iNPC/Neuronal-like. These patterns are broadly reminiscent of normal brain development, whereby early RG cells are the progenitors of both neuronal and glial lineages. Ordering the top differentially expressed genes by pseudotime, used here as a proxy for developmental-like progression rather than a confirmation of directionality (Fig. 3B), further suggested that cancer cells may recapitulate aspects of normal neurogenesis and gliogenesis. Early in pseudotime, we found that the genes *GFAP, VIM, CLU*, and *H3F3* – known to support progenitor and structural identity, survival, and active chromatin remodeling during early CNS development– were mainly expressed by RG-like cells, followed by GPC/AC-like cells.

**Figure 3.**
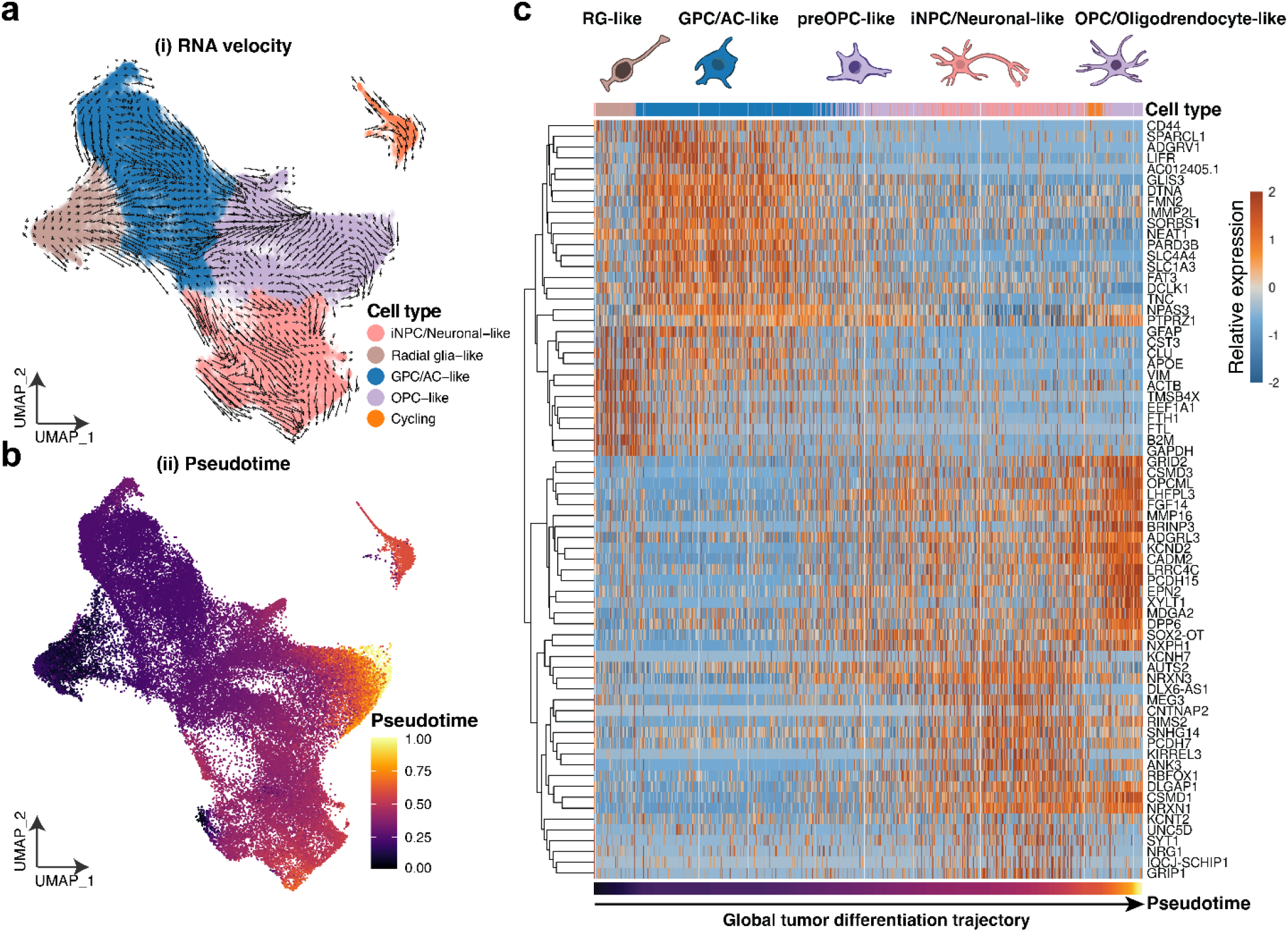
RNA velocity and pseudotime analysis reveal differentiation trajectories and cell state heterogeneity specific to IHGs. (a) RNA velocity vectors on UMAP embeddings of single-nuclei transcriptomes. (b) Corresponding pseudotime values inferred by the RNA velocity model, with lighter colors representing cells at later stages of the differentiation continuum. (c) Heatmap of differentially expressed genes along the tumor differentiation trajectory. Gene modules are ordered based on their expression patterns along the trajectory

At later pseudotime, genes such as *APOE, SPARCL1, CD44*, and *GLIS3* became more prominent and were mainly expressed in GPC/AC-like cells, reflecting a potential shift towards astrocytic-like differentiation^33,41^.

We can also note the expression of *ADGRV1, PTPRZ1, TNC, NEAT1*, and *SLC1A3* around the same pseudotime, genes of an astrocytic lineage that retain progenitor traits related to migration, extracellular matrix remodeling, and metabolic support typical of rapidly dividing cancer cells^42,43^.

At the latest pseudotime, we detected the expression of the synaptic receptor and adhesion genes *GRID2, ADGRL3*, together with *FGF14*, the tumor-suppressor gene *OPCML, PCDH15*, while mapping to OPC/Oligodendrocyte-like^39,44^. *NRG1*^45^, *NRXN3*^46^, and *RBFOX1*^47^, the voltage-gated potassium channel *KCNH7*^48^, were also found to be expressed at late pseudotime and enriched in iNPC/Neuronal-like cells. Finally, we found that *NRXN1*^46^, *DLGAP1*^49^, and *CSMD1*^50,51^ were expressed in both OPC-like and iNPC/Neuronal-like cells, illustrating an intermediate or uncommitted transcriptional state^39^.

Averaging single-cell RNA velocity by sample and cancer-cell state (Fig. S5D) showed the slowest velocity in RG-like cells (18 +/- 7.37), followed by GPC/AC-like (21.09 +/- 5.77), indicating that these cells undergo comparatively few transcriptional changes over time. In contrast, OPC-like (35.42 +/- 11.61) and iNPC/Neuronal-like (42.32 +/- 15.62) cells showed faster RNA velocity trends, suggesting active transcriptional changes potentially associated with differentiation, though functional validation would be needed to confirm these interpretations. Overall, our RNA velocity and pseudotime analyses suggest a developmental-like organization in which a subset of RG-like and GPC/AC-like cells retain stem-like properties that could drive a dysplastic development process resembling neuronal and glial-like lineages. We note that the velocity vectors serve as trend-level observations rather than definitive trajectory inference. Nevertheless, the data show a developmental-like hierarchy, in which each lineage acquires some degree of specialization without achieving complete differentiation.

### Cellular composition of IHGs vs other fusion-positive gliomas

Given the developmental-like hierarchy and underlying heterogeneity, we next asked whether the cellular composition of IHGs and other fusion-positive gliomas varied with age, fusion, and tumor type.

With most tumors containing all five cancer cell states (Fig. 1C, Fig. S1G), we calculated the lineage scores using the previously defined cellular states (OPC-like, RG-like, iNPC/Neuronal-like, and GPC/AC-like).Based on the lineage plot, we observed a higher proportion of iNPC/Neuronal-like cells in IHGs, and to a lesser extent, an increase in GPC/AC-like and RG-like cells (Fig. 4A, Fig. S5G; FDR < 0.001).

**Figure 4.**
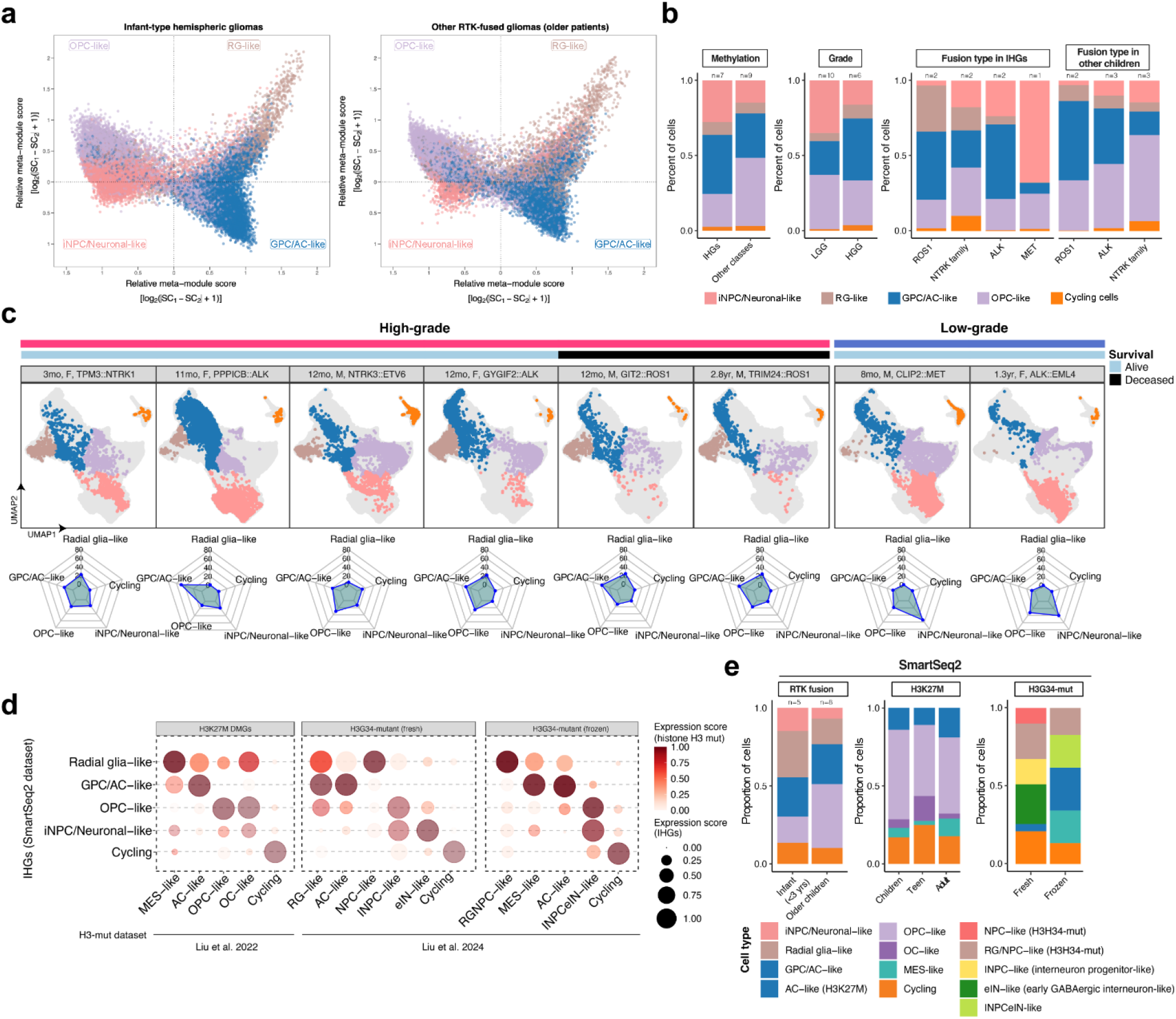
Cellular composition of IHGs across age, fusion, and tumor subtypes. (a) Two-dimensional projection of malignant cellular states. Each quadrant represents a distinct cellular state defined by meta-module activity. Individual malignant cells are plotted as dots, positioned according to their relative scores across the meta-modules. Dot colors indicate tumor cell population identities. (b) Composition of tumor cell populations grouped by methylation class, histological grade, and fusion subtype. (c) UMAP embeddings of single-nucleus transcriptome from IHG patients color-coded by cancer metaprogram and ordered by age. Radar plots showing the proportional distribution (%) of cancer cell metaprograms for each sample. (d) Bubble plots showing the projection of IHGs (SmartSeq2 dataset) tumor cell programs onto two H3-mutated high-grade glioma datasets: H3K27M and H3G34-mutant (Liu et al. (2022 and 2024, respectively)). (e) Composition of tumor cell populations (SmartSeq2 dataset) across tumor type (RTK-fused vs H3K27M vs H3G34-mutated).

When examining individual infant gliomas, all but one tumor had at least 10% RG-like cells (Fig. 4B,C and Fig. S5E,F). The near-universal presence of RG-like cells across IHGs is consistent with these tumors arising during a developmental window in which radial glia constitutes the brain’s dominant progenitor pool. This population may reflect both the cell of origin and a stem-like reservoir that sustains tumor growth — a feature likely enabled by the infant brain microenvironment and distinct from the cellular hierarchies observed in gliomas arising later in life. Except for ROS1-fused patients, IHGs and other fusion-positive gliomas with low-grade histology contained more iNPC/Neuronal-like cells (∼30%) relative to older children’s tumors (<10%), which instead were depleted in iNPC/Neuronal-like cells and enriched in OPC-like cells (FDR <0.001; Fig. 4B)

Moreover, we found that the four ROS1-fused tumors contained almost no iNPC/Neuronal-like cells in contrast to ALK- and NTRK-fused tumors (Fig. 4B). Our data hint at a fusion-specific developmental timing and/or effect: infant tumors are enriched for iNPC/Neuronal-like cells, suggesting an earlier oncogenic hit during development and/or increased plasticity. Notably, ROS1-fused tumors overall (IHG included) showed reduced representation of this lineage compared to tumors harboring other types of fusions (Fig 4B).

We next compared RTK-fused gliomas to Smart-seq2 datasets from H3K27M- and H3G34-mutated pediatric gliomas^7,11^ (Fig. 4D,E). Although distinct, RG-like cells aligned most closely to the MES-like population of H3K27M gliomas, and to both RG- and iNPC-like populations in G34-mutant tumors. GPC/AC-like cells were matched with AC- and MES-like cells in H3K27M tumors, whereas iNPC/Neuronal-like cells were associated with more mature interneuron populations in both H3-Mutant cohorts (Fig. 4D). These findings suggest both overlapping features and glioma subtype-specific differences among cancer cell states.

### The open chromatin landscape of IHGs and other fusion-positive pediatric gliomas

Given the transcriptional diversity of cancer cell populations in RTK-fused gliomas, we next asked whether this heterogeneity extends to the chromatin and gene-regulatory levels. We measured gene expression in the same cell with chromatin accessibility using joint snRNA/ATAC-seq (10X Multiome) in order to capture changes in transcription factor (TF) motif binding availability and cis-regulatory DNA elements (CREs), which are known to influence normal tissue and cancer developmen^52^. Focusing on the cancer cells, we characterized n=68’024 epigenomes in 15 patients and direct annotation transfer from snRNA-seq (Fig. 2A) confirmed that the chromatin landscape is both patient-specific and heterogeneous (Fig. 5A, S6 A-C).

**Figure 5.**
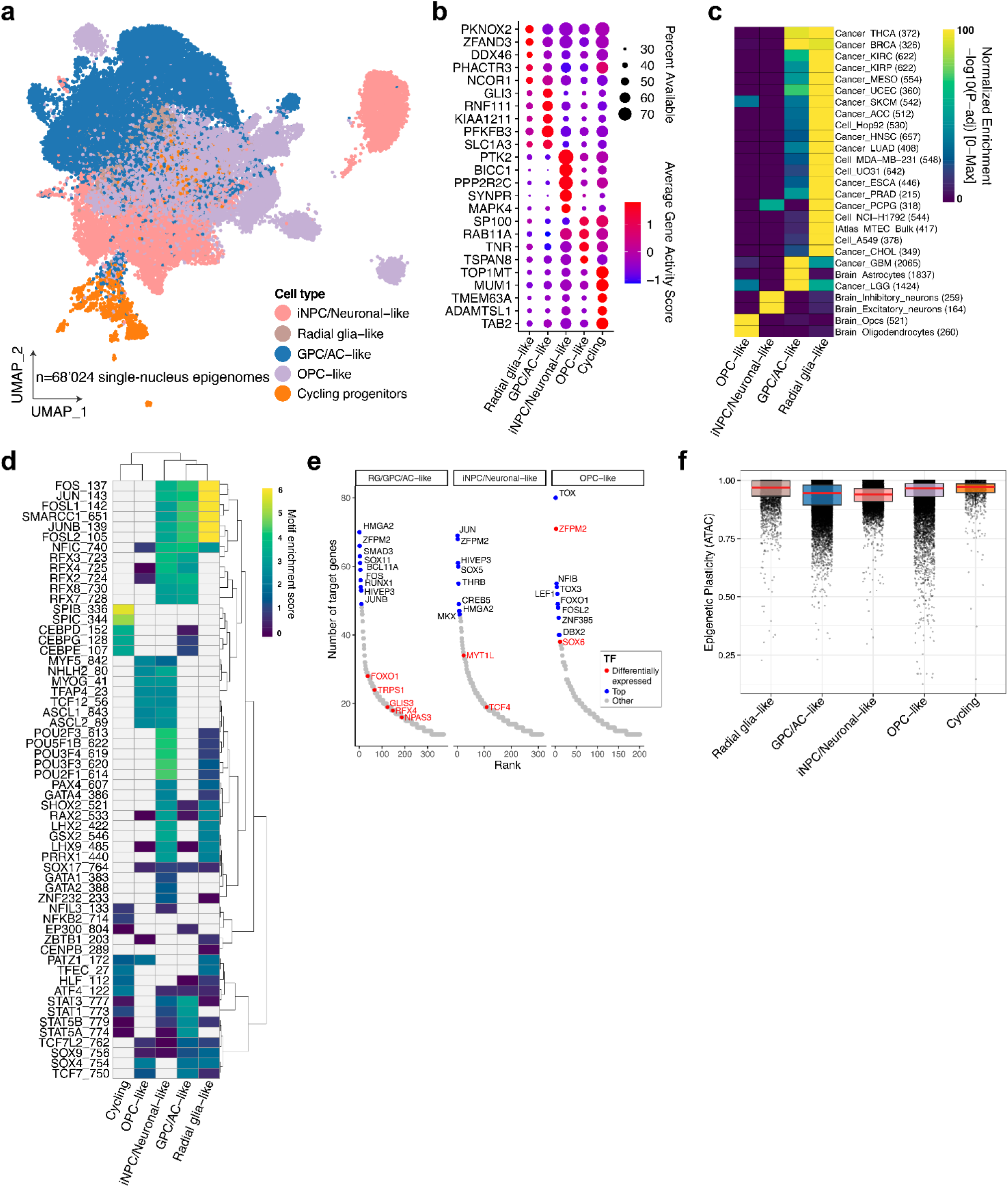
Epigenetic landscape of RTK-fused gliomas. (a) UMAP of single-nucleus ATAC-seq data from tumor cells after Harmony batch correction and colored by cancer cell type as defined by snRNA-seq. (b) Dot plot showing of top genes with differential gene activity scores (prediction of gene expression based on the accessibility of the chromatin, i.e. number of fragments detected in the vicinity of the gene). Genes that are highly accessible have a high gene score. Dot size as percent of cells with gene accessible. (c) Chromatin accessibility feature enrichment analysis using a curated bulk ATAC-seq dataset; heatmap shows enrichment scores between cancer cell populations and bulk ATAC-seq experiments. (d) Motif enrichment analysis on differential peaks across cancer cell populations. Grey cells indicate motifs that were not detected for a given cell type. (e) Top transcription factor (TFs) modules (regulons) derived by inferring gene regulatory networks (GRNs) using Pando. Regulons we derived per cancer cell type, then ranked together according to the number of target genes. Top regulons and regulons for which the gene was differentially expressed across populations colored in blue and red, respectively. (f) Boxplots showing the distribution of per-cell plasticity scores (0 = regulatory commitment, 1 = maximal epigenetic plasticity) based on chromatin accessibility and RNA-derived cell types. Plasticity was computed as the normalized Shannon entropy of softmax-transformed module scores, reflecting the degree to which each cell simultaneously engages multiple regulatory programs.

Gene activity scoring, assessed through chromatin accessibility at gene body and promoter regions, revealed cell type-specific chromatin signatures (Fig. 5B): RG-like cells scored for TFs *PKNOX2* and *NCOR1*, both linked to glioblastoma and neural stem cells maintenance^53,54^. GPC/AC-like cells showed *GLI3* enrichment, a mediator of Shh signaling that can have a role in cortical development^55^, while iNPC/Neuronal-like cells featured synaptic protein *SYNPR*^56,57^, along with *PTK2* (*FAK*) and *MAPK4*, two regulators of cell survival. OPC-like cells were marked by the oligodendrocyte marker *TNR*. We then generated pseudo-bulk replicates of each cancer cell populations and compared them to annotated bulk ATAC-seq datasets^29,58^ of normal and cancer tissues (Fig. 5C). The chromatin state of OPC-like, iNPC/Neuronal-like, and GPC/AC-like shared features associated with their wild-type counterparts. In contrast, features of RG-like and GPC/AC-like cells overlap with other cancers such as glioblastoma and low-grade glioma, and breast and thyroid cancers (Fig. 5C). These overlapping profiles further suggest that RG-like and GPC/AC-like cells lack lineage commitment and may harbor stem-like cancer properties.

Motif enrichment analysis identified brain and lineage-defining TF motifs involved in cell cycle regulation, survival, development, and oncogenic processes (Fig. 5D), but their distribution across the cancer cell states lacked lineage-specific patterns. For instance, motifs for the *FOS/JUN* family and the chromatin remodeling complex *SMARCC1* were enriched in most populations. Neural developmental motifs from the *RFX, POU*, and *TCF* families were also detected: *POU2F1* (*OCT1*) and *POU3F2* were enriched in iNPC/Neuronal-like cells and, to a lesser extent, RG-like cells. Motifs *RFX2-4 and RFX7-8* were also enriched in GPC/AC-like cells^59,60^, *TCF7* and *TCF12* in OPC-like cells^61^, and *SOX4, SOX9*, and *SOX17* to a lesser extent in several populations^62^. Interestingly, myogenic TFs *MYOG* and *MYF5* were also detected in OPC-like and iNPC/Neuronal-like cells.

Although RTK-fused gliomas lack a defined MES-like transcriptomic state (Fig. 1 B,C), these motifs may reflect the potential to engage mesenchymal regulatory programs^7,36,63,64^.

By correlating chromatin-accessibility peaks (snATACseq) with gene expression (snRNAseq) within the same cell, we identified 46’615 significant peak-to-gene links, thereby assigning candidate CREs to their putative target genes in RG/GPC/AC-like, iNPC/Neuronal-like, and OPC-like (n=3 IHGs, Fig. S6E). We then applied Pando^55^ to each of these populations to independently infer gene regulatory networks (GRNs) and obtained 18’915 regulons (Table S4, Fig. 5E). Ranking regulons by number of target genes revealed top TFs including *HMGA2, SOX11*, and *BCL11A* in RG/GPC/AC-like cells, *SOX5* in iNPC/Neuronal-like cells, and *ZFPM2* (*FOG2), NFIB*, and *SOX6* in OPC-like cells. Again, inferred regulons deviated from canonical neurodevelopmental GRNs. For instance, *SOX11*, though traditionally linked to neuronal lineage commitment, was also identified in RG/GPC/AC-like cells. In iNPC/Neuronal-like cells, *SOX5* emerged as top regulon, which together with *MYT1L* and *TCF4* reinforces a pro-neuronal, glioblastoma program, aligning with its established role in enforcing neuronal identity ^65-67^. Top regulons in OPC-like cells included *DBX2, ZFPM2 (FOG2)*, and *NFIB* with known roles in both astrocytic and neuronal differentiation. Finally, *FOS, FOSL2, JUN*, and *JUNB* emerged as regulators spanning all cancer cell populations, especially in RG- and GPC/AC-like cells, reflecting RTK-driven MAPK dysregulation.

We calculated an epigenetic plasticity score for the five cancer cell populations based on the entropy of their chromatin accessibility across gene activity scores. All populations displayed uniformly high plasticity (Fig. 5F), indicating that cancer cells are not committed to regulatory lineage programs but rather occupy multiple epigenetic states simultaneously.

Taken together, multi-omic profiling revealed cell-type-specific CREs and GRNs that corroborate the extensive heterogeneity observed at the transcriptomic level. Open chromatin regions in RG-like and GPC/AC-like cells resembled those in other cancers and retained brain-specific features. Overlapping TF motifs and regulons between cancer cells and neurodevelopmental programs underscore their regulatory plasticity. Notably, FOS/JUN GRNs detected in all populations point to RTK/MAPK dysregulation as a core oncogenic program promoting proliferation and survival.

### Defining the myeloid landscape in infant RTK-fused gliomas

While glioma cell composition varies across brain regions and developmental stages, the identity and function of tumor-associated myeloid cells in IHGs and other RTK fusion-positive gliomas remain largely unexplored. Our atlas identified a prominent myeloid compartment that accounted for 22% of all cells (Fig. 1B) and was detected in every sample profiled by 10x Multiome (SS2 samples not included, as pre-processing involved CD45^+^-cells depletion). Reintegration of all myeloid cells revealed seven transcriptionally distinct subpopulations (Fig. 6A), which we collectively refer to as tumor-associated macrophages (TAMs), as they encompass both resident microglia (MG) and infiltrating myeloid-derived macrophages (MDMs) (Fig. S7). Clusters were annotated based on differentially expressed genes, enriched GO biological processes, and GSEA (Fig. 6 B,C).

**Figure 6.**
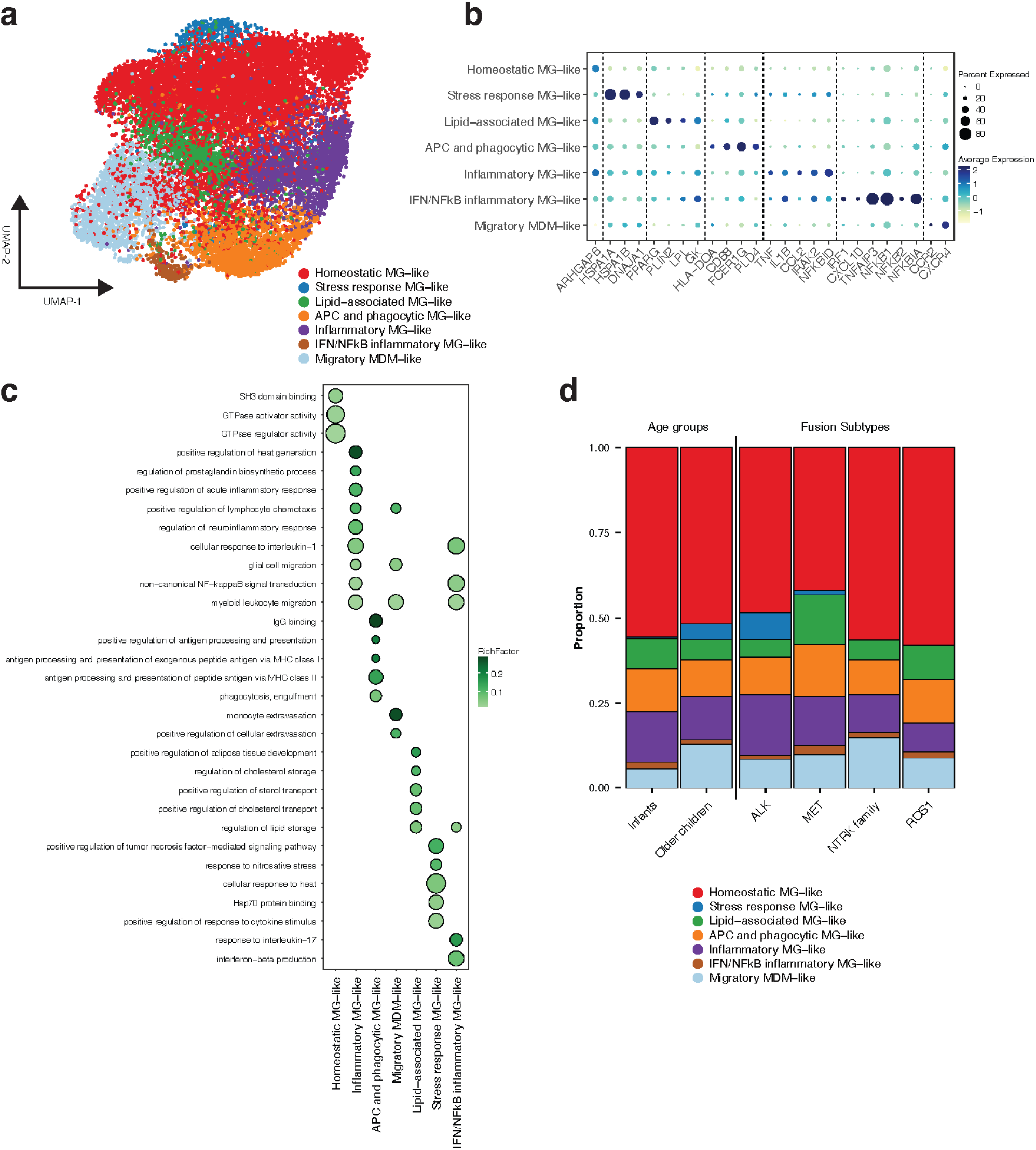
The myeloid landscape of RTK-fused gliomas. (a) UMAP projection of myeloid single cells colored by subtype. (b) Dot plot of average expression and percent expressed in cluster for selected genes differentially expressed across myeloid subpopulations. (c) Dot plot of gene ontology (GO) terms, showing gene ratio scores for each term (Rich Factor) across different myeloid subpopulations. (d) Composition of myeloid subpopulations grouped by age and fusion subtype

The TAM subpopulations included a homeostatic microglia (MG)-like state, stress-response MG-like, lipid-associated MG-like, antigen-presenting cell (APC) and phagocytic MG-like, inflammatory MG-like and a subset of IFN/NFkB inflammatory MG-like cells, as well as a migratory MDM-like TAM subpopulation. The homeostatic state is marked by the expression of GTPase-associated genes linked to a non-inflammatory MG phenotype^68^ and negative GSEA for inflammatory pathways. The stress-response MG-like subpopulation is characterized by increased expression of stress-response pathways and heat shock proteins (*HSPA1A, HSPA1B, DNAJA1*). The lipid-associated MG-like state shows upregulation of pathways related to adipose tissue development and cholesterol homeostasis, with expression of several lipid metabolism genes (e.g., *PPARG, LPL, PLIN2, GK*). The APC and phagocytic MG-like cluster is enriched for gene expression programs and transcripts related to antigen processing (*HLA-DOA*) and phagocytosis (*CD68, FCER1G, PLD4*).

Within the TAMs with inflammatory signatures, we distinguish 2 subsets: inflammatory MG-like cells, that upregulate several inflammatory pathways, and IFN/NFκB inflammatory MG-like cells, which, in addition, show an upregulation of genes related to signaling via NF-κB. (e.g. *NFKB1*). Lastly, a migratory MDM-like state is marked by the expression of MDM genes and an upregulation of genes (e.g., *CCR2, CXCR4*) and GO terms related to cell migration.

We observe some variation in the relative abundance of these subpopulations with patient age (Fig. 6D). The most prominent trend was an increase in the proportion of the stress-response MG-like and migratory MDM-like cells in older children relative to infants. These findings are consistent with the more inflammatory myeloid landscapes and higher abundance of MDM states described in adult HGG^69,70^. Comparison across fusion subtypes was limited due to small numbers, but we observed a near absence of stress-response MG-like cells in NTRK- and ROS1-fused gliomas (Fig. 6D).

Together, these functional profiles reveal an abundant myeloid compartment comprising distinct TAM states with potential roles in tumor biology, whose relative abundance appears to be shaped by developmental stage and, to a lesser extent, oncogenic driver.

### The spatial architecture of IHGs

Gliomas can exhibit diverse spatial patterns, ranging from disorganized cell distributions to well-defined niches characterized by structured interactions between malignant and non-tumorous cells^71^. To map these architectures in IHGs, we applied MERFISH and Xenium spatial transcriptomics to 8 tumor sections. MERFISH was used to characterize four samples from our main cohort (Fig. 1A), spanning four fusion subtypes *PPP1CB::ALK, CLIP2::MET, GIGYF2::ALK, GIT2::ROS1* (Figure 7A-B, Figure S8). The MERFISH panel comprised 500 mRNA targets (Fig. 7A-B), including 378 transcripts corresponding to the 5 major malignant cell populations identified in our atlas (Fig. 2A) and 122 pan-cancer and immune markers (Table S5). Four additional tumor sections were profiled using the Xenium panel comprising 311 targets: 261 genes from the 10x Genomics human brain base panel and an additional 50 custom genes selected to mark common metaprograms in high-grade gliomas. DAPI-based and Baysor^72,73^ cell segmentation, followed by label transfer from our snRNA-seq atlas (Fig. 1B), resolved approximately 1 million cells whose proportions closely mirrored snRNA-seq data (Fig. 7B,C; Fig. S8A,B).

**Figure 7.**
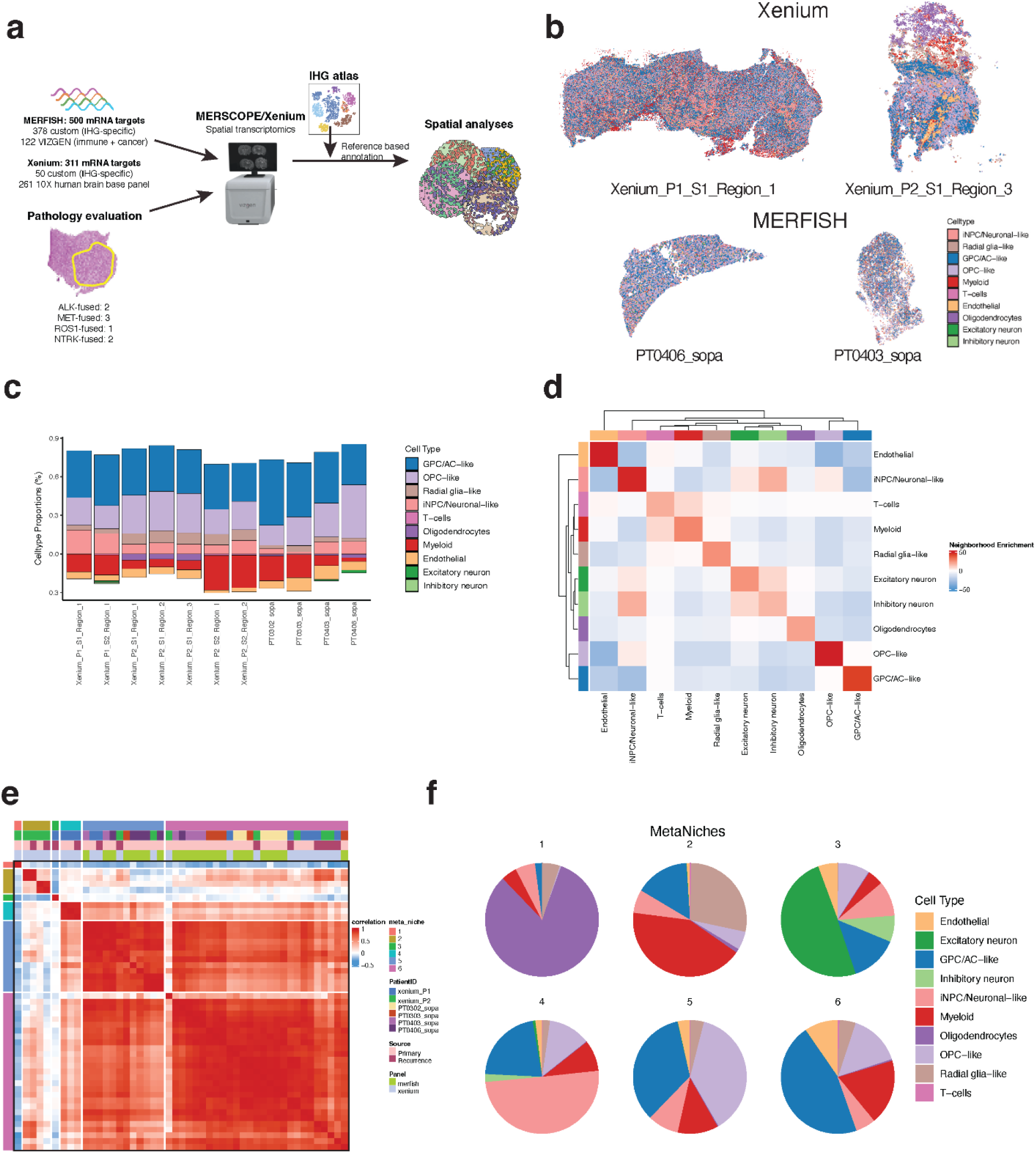
The spatial architecture of RTK-fused gliomas. (a) Schematic of the spatial transcriptomics workflow in 8 samples using Xenium (n=4) and MERFISH (n=4). (b) Representative images of each IHG section processed with either Xenium (top) and MERFISH (bottom), with cells colored by cell type. (c) Cell type proportions across sections. (d) Neighborhood enrichment analysis (average z-score values). (e) Heatmap displaying correlation of cell type proportions for each niche across sections, clustered into six “metaniches.” (f) Cell type proportions within each metaniche

Spatially, most tumor sections exhibited a largely disorganized distribution of cell types (Fig. 7B). However, heterogeneity was evident between patients and samples, with some sections showing areas enriched for specific cell populations. To characterize the spatial organization of IHG tumor sections, we performed neighborhood enrichment analysis across all samples (Fig. 7D). T-cells and myeloid cells exhibited self-enrichment, as did excitatory and inhibitory neurons, consistent with known immune clustering and laminar neuronal organization. By contrast, malignant metaprograms showed no consistent co-occurrence patterns, suggesting spatially variable rather than structured malignant niches. We next identified spatial niches within each sample and clustered them across all samples using pairwise correlation, yielding 6 recurrent spatial “metaniches” (Fig. 7E,F). Two metaniches were predominantly composed of non-tumoral cell types (metaniches 1, enriched for oligodendrocytes, and metaniche 3, enriched for excitatory neurons). Metaniche 2 encompassed the highest proportions of myeloid and radial glial like cells, but was restricted to 2 tumor sections. The remaining three metaniches were dominated by malignant metaprograms: metaniche 4 is iNPC/neuronal-like enriched, metaniche 5 contains mostly OPC-like and GPC/AC-like cells, while metaniche 6 contains predominantly GPC/AC-like cells. Apart from the GPC/AC-like- and OPC-like-dominated metaniches, the remaining metaniches are mostly sample-specific, highlighting the heterogeneity in the spatial organization of these tumors.

Our spatial analyses revealed that tumors appear spatially disorganized but can harbor discrete cancer cell-specific spatial clusters with high heterogeneity between tumors.

## DISCUSSION

Pediatric tumors are increasingly understood as organ-like systems, where cancer cells retain features of and interact with the developing tissue from which they originate. At the heart of this view is the increasing recognition that many tumors contain diverse cellular states, of which a subset of cancer cells drives tumor growth and diversification. These cells give rise to a range of malignant progenitor-like and variably (un)differentiated cell types, resembling the structured lineage progression observed in healthy tissues.

By profiling a rare cohort of pediatric gliomas driven by ALK/ROS1/MET/NTRK fusions, including IHGs, we identified five distinct cancer cell populations that have features reminiscent of their tissue of origin, while also exhibiting transcriptional programs tied to their oncogenic drivers. By modeling cell trajectories, we propose that RG-like and GPC/AC-like cells constitute a pool of cancer progenitors with stem-like features and the ability to adopt neuronal, astroglial, and oligodendrocytic gene expression programs, resulting in heterogeneous and dysplastic cell populations.

The CNS is composed of a large variety of cell types, many of which are transient and strictly regulated in time and space. During prenatal human cortical development, extensive proliferative compartments of radial glial cells with neurogenic capacity generate intermediate precursor cells and excitatory neurons. Gliogenesis follows shortly after the start of neurogenesis, with multipotent glial precursor cells further differentiating along the astrocytic and oligodendrocytic lineages. In neonates and young children who present with IHGs shortly after birth, it is conceivable that the oncogenic event takes place in early neural progenitors. Notably, RG-like populations were detected in all IHGs, and iNPC/Neuronal-like populations were enriched in IHGs driven by ALK and NTRK fusions but were sparse or absent in ROS1-driven IHGs and RTK-fused tumors in older patients. This suggests that oncogenic transformation may occur within different progenitor compartments, e.g. multipotent glial precursor cells lacking neurogenic potential (Figure 8). Alternatively, the oncogenic drivers themselves may preferentially promote the expansion of specific populations. This and other studies^74,75^ suggest that *ROS1*-fused IHGs occupy a distinct position within infant gliomas and may warrant separate biological and clinical consideration.

**Figure 8.**
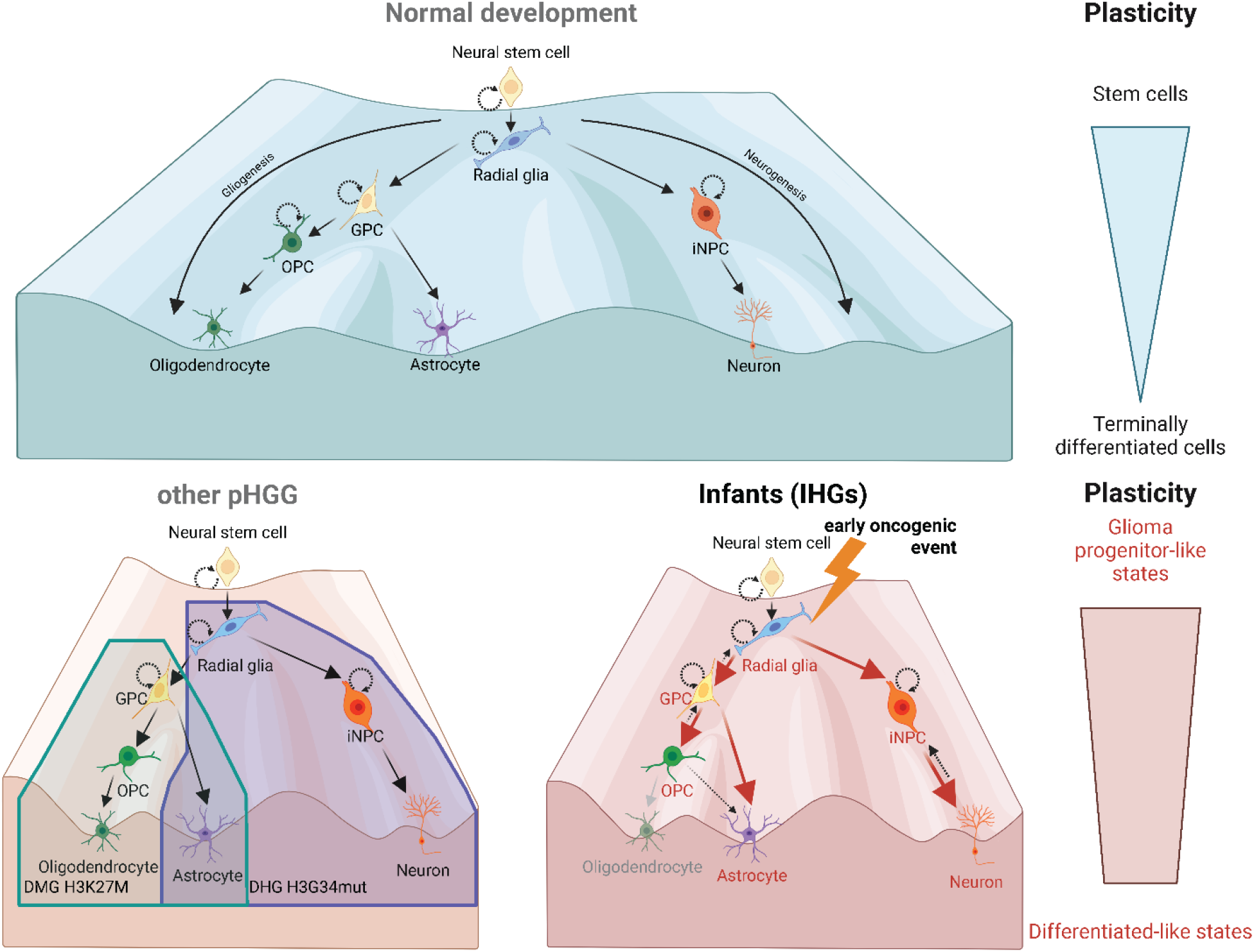
Proposed model of IHG tumor formation.

We report here the cellular composition of IHGs and other gliomas with RTK fusions, distinct from previously published pediatric and adult gliomas. Adult gliomas, for instance, frequently comprise cell populations with a mesenchymal-like signature^7,36^, which is largely absent in IHG, consistent with previous findings suggesting that mesenchymal features in gliomas become more prominent with age^7,36^. Although RTK-fused gliomas do not display a MES-like transcriptomic state, the presence of *MYOG* and *MYF5* motifs indicates that a mesenchymal regulatory program could be epigenetically primed. This ‘latent MES potential’ aligns with the high plasticity we observe across chromatin states and may become transcriptionally engaged under specific microenvironmental conditions, such as hypoxia or therapeutic pressure^64^. Conversely, the presence of RG-like compartments in IHG, absent in adult gliomas^64^, further supports the concept of divergent developmental origins. Alongside single-cell studies of pediatric HGGs driven by histone mutations, our study further underlines the diversity in cellular landscapes in pediatric gliomas. While H3G34-mutant tumors arise from disruptions in interneuron lineage development^11^ and H3K27-altered DMGs stem from OPC-like progenitor cells^7,8^, our study shows that IHGs exhibit a developmental trajectory from RG-like cells towards iNPC/Neuronal-like, GPC/AC-like, and OPC-like lineages. Notably, these patterns imply distinct developmental origins, either shaped by the type of alteration or the timing of the oncogenic hit.

We propose that oncogenic fusions drive transformation and expansion of progenitor-like cancer cells, which retain both high proliferative capacity and the ability to engage neuronal and/or glial gene expression programs. This high plasticity, supported by integration of single-cell RNA-seq and ATAC-seq data, underlies a dynamic and diverse cancer cell composition. This, in turn, can explain the striking histological heterogeneity observed in IHGs, which spans the LGG to HGG spectrum. This diversity aligns with the wide range of clinical outcomes, from rapidly progressive disease with early mortality to cases that resolve with minimal or no post-surgical treatment^3^.

Our study supports the widely accepted hypothesis of developmental origins of pediatric brain tumors; however, we expect that future studies exploring interpatient heterogeneity will reveal biologically relevant subpopulations, expanding on these first models of pediatric glioma organization. Further, annotating cells that exist on a continuum of states, rather than as discrete populations, has important limitations, and we observe overlapping transcriptomic and chromatin accessibility features within malignant cells, mainly between RG- and GPC/AC-like cells. Chromatin accessibility analyses revealed increased activity of transcriptional effectors from the STAT3 and MAPK pathways, promoting cell proliferation and survival. Oncofusions involving these RTKs are not unique to gliomas^76,77^. Their ability to transform cells across multiple developmental contexts and tissues likely contributes to the clinical and histopathological heterogeneity. Moreover, this pathway dependency could explain why these patients respond well to targeted RTK inhibitors ^78^ and have a better prognosis, in contrast to the treatment resistance observed in all other pHGGs.

We identified a myeloid compartment comprising seven subpopulations of TAMs. Transcriptionally, these spanned a broad spectrum of biological programs — from homeostatic and metabolically active states to pro-inflammatory and lipid-associated programs. Whether these differences contribute to the clinical behavior of IHGs, and whether specific TAM subpopulations occupy preferential spatial niches relative to malignant cells, remains unclear^79,80^. Spatial transcriptomic analyses revealed overall unstructured and heterogeneous tissue architectures.

Understanding these spatial niches will need to be the subject of future work, and may uncover therapeutic opportunities to disrupt microenvironments that sustain tumor growth^71,79,80^.

Several questions remain. Firstly, does the LGG-to-HGG continuum in IHGs simply reflect the developmental window in which an RTK fusion arises? While we propose that the temporal context is important, additional factors, such as type of fusion, co-occurring genetic alterations or epigenetic modifiers, may also drive tumor cell composition. Moreover, although our data support a hierarchical organization reminiscent of early neural development, it remains unclear whether individual tumor cells can transition between cell states or undergo “multidirectional” de- or trans-differentiation. External cues and “fate switching” from the tumor microenvironment may further shape this plasticity. Rigorous *in vitro* and *in vivo* models will be essential to test these possibilities. Lastly, the transcriptional and functional overlap among cancer cell populations from different pediatric gliomas is yet unknown. Our atlas provides an important snRNAq/ATACseq IHG-specific reference, upon which future cross-entity comparisons can be built once appropriately matched cohorts become available.

In conclusion, our study resolves the cellular diversity of IHGs and other RTK-fused gliomas, identifying cancer cell states resembling early neural development. Modeling oncogenic events in a neural developmental context, experimentally tracing cell trajectories using murine or organoid systems, will be essential to determine how RTK fusion timing and cellular context shape IHG development and cellular composition. Such models will also be instrumental for developing therapeutic strategies, ultimately contributing to the discovery of more effective treatments for the most vulnerable patients, including infants.

## METHODS

### Clinical cohort and data collection

All patient material and genomic data used in this study were obtained with proper informed consent obtained at the University Children’s Hospital Zürich, The Hospital For Sick Children in Toronto, the University Hospital Motol in Prague, the Children’s Hospital of Philadelphia, and Boston’s Children’s Hospital/Dana-Farber Cancer Institute.

### Tissue collection and processing

Tumor specimens were collected from infants diagnosed with hemispheric gliomas between the years 2012 and 2022 following surgical resection. The tissues were flash frozen in OCT following surgical resection and biobanked as appropriate.

For snRNA/ATAC-seq, frozen tissue pieces were weighed on dry ice and transferred to a petri dish. On ice, 500 μL of lysis buffer (composed of Lysis Reagent, Reducing Agent B, and Surfactant A from 10x Genomics) was added, and the tissue was chopped with scissors. The tissue was then transferred to a 2 mL glass douncer and dissociated with 8 strokes using the loose pestle, with a total lysis time of 5 minutes. After lysis, 500 μL of the lysate was passed through a Nuclei Isolation Column (10x Genomics) and centrifuged at 16,000 rcf for 20 seconds at 4°C to remove debris. The sample was briefly vortexed (10 seconds) and an aliquot was taken to count nuclei using a 1:1 dilution with AOPI. The count was used to determine the volume needed for resuspension. The sample was then centrifuged at 500 rcf for 3 minutes at 4°C; the supernatant was removed, and the nuclei pellet was resuspended in Nuclei Separation buffer (comprising Nuclei Extraction buffer, PBS, BSA, and RNase Inhibitor). Miltenyi Anti-Nucleus beads (50 μL per million nuclei) were added, and the mixture was incubated on ice for 15 minutes. An additional 2 mL of Nuclei Separation buffer was added before passing the sample through an LS column attached to a MACS MultiStand with a QuadroMACS separator. The column was washed twice with 1 mL of buffer, then removed and flushed with 1 mL of buffer into a collection tube. The purified nuclei were pelleted (500 rcf for 5 minutes), washed with Wash Buffer (1× PBS, 10% BSA, and RNase Inhibitor), and recounted using AOPI on a haemocytometer.

For scRNA-seq (Smart-seq2), whole cells isolated from tumor tissue acquired at the time of surgery were immediately processed by mechanical as well as enzymatic papain-based dissociation for 30 min at 37°C according to the Brain Tumor Dissociation Kit (Miltenyi Biotec) protocol. Single-cell suspensions were filtered through a 70 μm strainer, centrifuged at 300 x *g* for 10 min, and re-suspended in phosphate-buffered saline (PBS) supplemented with 1% bovine serum albumin (BSA) for subsequent fluorescence-activated cell sorting (FACS). Single-cell suspensions captured from fresh tumor tissue and re-suspended in PBS + 1% BSA were stained with 0.5 μM calcein AM (Life Technologies, C3100MP) and 0.33 μM TO-PRO3 iodide (Life Technologies, T3605) for 15 min at room temperature and subsequently placed on ice. Single-cell sorting was performed on a SH800 sorter (Sony) using 488 nm (calcein AM, 530/30 emission filter) and 633 nm (TO-PRO-3, 665/30 emission filter) lasers. Unstained and single-stained controls were included for all tumors. Viable cells were identified by positive staining for calcein AM and negative staining for TO-PRO-3. Doublet discrimination was performed by stringent singlet-gating based on back scatter area (BSC-A) versus back scatter width (BSC-W) setting. Single CD45-viable tumor cells were sorted into 96-well plates containing cold TCL buffer (Qiagen, 1031576), briefly spun down, snap frozen on dry ice, and then stored at -80°C. Whole transcriptome amplification, library preparation and sequencing of single cells/nuclei were performed using the Smart-seq2 modified protocol as described previously (Liu et al 2024). RNA was purified with Agencourt RNAClean XP beads (Beckman Coulter). Oligo-dT primed reverse transcription (RT) was performed using Maxima H Minus reverse transcriptase (Life Technologies) and a template-switching oligonucleotide (TSO; Qiagen). PCR amplification (20 cycles) was performed using KAPA HiFi HotStart ReadyMix (KAPA Biosystems), followed by Agencourt AMPure XP bead (Beckman Coulter) purification. Libraries were generated using the Nextera XT Library Prep kit (Illumina). Libraries from 768 cells with unique barcodes were combined and sequenced using a NextSeq 500/550 High Output Kit v2.5 (Illumina).

### Single-nucleus RNA/ATAC-seq library preparation

Nuclei were loaded onto a 10x 3’ GEM X chip targeting 6’000 cells to avoid doublets. After droplet generation, samples were transferred to pre-chilled strip tubes and incubated in a thermocycler. cDNA was recovered using the 10x Recovery Agent, cleaned with a Silane DynaBead mix, and amplified for 11 cycles. Libraries were further purified using SPRIselect beads, and quality was assessed with a Bioanalyzer to determine the cDNA concentration. Final library preparation followed the Chromium GEM-X Single Cell 3’ Reagent Kits v4 protocol, with PCR cycle modifications based on the calculated cDNA concentration.

The molarity of each 10x library was calculated based on library size as measured bioanalyzer (Agilent Technologies) and qPCR amplification data (Roche, cat. no. 07960140001). Gene Expression libraries were sequenced on Illumina’s NovaSeqX with the following run parameters: read 1 – 28 cycles, read 2 – 90 cycles, index 1 – 10 cycles and index 2 – 10 cycles.

### Bulk RNAseq

Total RNA samples were quantified by qubit RNA kit (Life Technologies) and quality assessed by Agilent Bioananlyzer. Libraries were prepared with 100 ng of total RNA using Stranded Total RNA prep with Ribo-Zero Plus kit (Illumina) with 13 cycles of amplification used. Final cDNA libraries were size validated using Agilent Bioanalyzer or Tapestation and concentration validated by qPCR (Kapa Biosystems/Roche). All libraries were normalized and pooled together, denatured with 0.2N NaOH and diluted to a final concentration of 260 pM. Pooled libraries were loaded onto an Illumina Novaseq V1.5 cartridge for cluster generation and sequencing on an Illumina Novaseq 6000 instrument (Illumina) Paired-end 101 bp protocol to achieve ∼40 million reads per sample.

### Methylation array and classification

For a subset of patients where methylation data was unavailable, genomic DNA was isolated from using standard extraction protocol. Briefly, 500 ng from genomic DNA samples were bisulfite converted using EZ DNA methylation kit (Zymo Research cat# D5001 or D5002) following manufacturer’s protocol with the following modification, samples were incubated in thermocycler for 16 cycles at 95°C for 30 seconds, 50 °C for 60 min, hold at 4°C. Eight microlitres from each sample were amplified following the Illumina Infinium HD Methylation assay protocol. The amplified DNA generated from these samples was fragmented, precipitated and 26ul from each of the samples was used to hybridize to an Infinium Human Methylation EPIC V2.0 beadchip. The beadchip was incubated at 48°C for 18 hours, then washed and single base extended as described in the Illumina protocol and scanned on the iScan, quantified in GenomeStudio (v.2011.1) (Illumina). The raw fluorescence intensities from each probe on the array was conferred into binary IDAT files for downstream analysis.

The Heidelberg Epignostic CNS Tumor Classifier (v.12.8) (formerly DKFZ brain classifier v.12.8)^81^ was used to classify the tumors from the IDAT raw files into one of 184 tumor classes, consistent with the 2021 WHO classification of CNS tumors^17^. Cases for which the classifier could not find a class were labeled a “*nonclassifiable*”.

### Cryosectioning and dehydration of the tissue for MERFISH

Each OCT block containing a fresh-frozen sample was treated separately. The block was transferred from an ultra-low temperature freezer at -80°C to a Leica CM3050S freezing microtome chamber at -20°C and allowed to sit for 30min. The tissue was oriented as parallel to the blade and trimmed in the cryostat until a complete section was obtained. Next, a first section was cut at 10µm and a MERSCOPE slide was lowered onto the flattened sample, centered. The slide was then transferred to a Petri dish placed in the chamber at -20°C. A second adjacent section was processed similarly. After 20min in the cryostat chamber, both slides were transferred under a chemical hood and incubated within a minute in 4% PFA for 15min at RT, prior to washing and incubating in 70% EtOH overnight at 4°C. One slide was used for verification of RNA integrity in situ, and the other was used for MERFISH.

## MERFISH

MERFISH was performed using MERSCOPE (Vizgen) and strictly according to the manufacturer’s protocol (“Fresh and Fixed Frozen Tissue Sample Preparation 91600002 Rev C”). Optional steps, including autofluorescence quenching and protein staining, were omitted. A clearing method for non-resistant fresh frozen tissue was used. Mounted MERSCOPE slides were processed within 10 days of cutting, and regions of interest were manually delineated, leaving about a 1 mm margin from the tissue. Segmentation was performed using Cellpose^72^ on cell boundary stain 1.

### Fusion Detection from bulk and Smart-seq2 RNA-seq

For both bulk RNA-seq and Smart-seq2 single-cell RNA-seq datasets, we used a custom pipeline that combined two fusion callers for accuracy^82,83^: STAR-Fusion (v. 1.13.0) and Arriba (v. 2.3.0). For Smar-seq2, FASTQ files from all individual cells were concatenated together. FASTQ reads were then aligned to the GRCh38 reference genome using STAR (v.2.7.11a) (passing recommended parameters for fusion detection, such as --chimSegmentMin 12 -- chimJunctionOverhangMin 12 --chimOutType WithinBAM Junctions). The resulting alignment outputs and chimeric junction files were then passed to STAR-Fusion to identify fusion transcripts. In parallel, we used Arriba with the STAR-generated Aligned.out.bam file to cross-validate fusions. Arriba was supplied with a blacklist (blacklist_hg38_GRCh38_v2.1.0.tsv.gz), known fusion references (known_fusions_hg38_GRCh38_v2.1.0.tsv.gz), and protein domain annotation (protein_domains_hg38_GRCh38_v2.1.0.gff3) to produce the fusion report.

For the Smart-seq2 samples, fused reads IDs from the STAR_Fusion and arriba reports for fusion of interest (*ALK, MET, ROS1, NTRK* family) we then mapped back to their original single cell. Fusion positive cells and well as the number of supporting fusion reads were added to the single-cell (Seurat) metadata object.

### Sn-RNA-seq data processing

Sequencing reads generated from the 10x Genomics Chromium platform were processed using Cell Ranger ARC (v.2.0.0) and aligned using the GRCh38 genome reference to generate both an ATAC fragment file and gene expression (RNA) count matrix. SoupX^84^ (v1.6.0) was used to remove ambient RNA contamination signal from the gene expression count matrices. Corrected gene expression matrices were then imported into Seurat^85^ (v.4.3.0) for downstream processing. First, cells expressing n<3 genes and genes expressed in n<3 cells were removed. Given that mitochondrial transcripts should not be found in the nucleus, mitochondrial genes were also removed from the gene expression matrix. In addition, low quality nuclei were filtered out based on the number of detected genes and transcripts, with nFeature_RNA > 0 and < 5,000, and nCount_RNA > 1,000 and < 20,000 for most samples. Cell cycle scores were calculated using cell cycle-specific gene sets provided by Seurat and the ‘CellCycleScoring’ function. Next, normalization and variance stabilization were performed using the SCTransform (SCT) function in Seurat, regressing out nFeature_RNA, nCount_RNA and cell cycle scores. Doublets were identified from normalized data using the DoubletFinder^86^ (v.2.0) package. Highly variable features were identified using the ‘FindVariableFeatures’ function with the “vst” method, selecting 3,000 genes. Principal component analysis (PCA) was conduction on these variable features, from which the cells were clustered into metaprograms using the ‘FindNeighbors’ and ‘FindClusters’ functions with a resolution parameter of 0.2 and the using the first 15 principal components. Finally, Uniform Manifold Approximation and Projection (UMAP) was used to visualize the cells.

### Cell type annotation and data integration

Prior to data integration, each sample underwent independent processing and cluster annotation. Differential expression analysis was performed on a per-sample basis using Seurat’s FindAllMarkers function, with parameters set to a minimum percentage expression threshold (min.pct) of 0.5 and a log fold change threshold (logfc.threshold) of 0.5. For each sample, the top differentially expressed genes for each cluster were identified and used for manual annotation. Cluster identities were assigned by cross-referencing these marker genes with established literature and publicly available datasets, including The Human Protein Atlas single-cell of the brain^87^ and the Spatio-Temporal Cell Atlas of the Human Brain (STAB^88^) datasets. Data integration was performed using the Reciprocal PCA (RPCA) method implemented in Seurat and using the SCT normalization method to correct for batch effects. Integration features were selected using the top 5000 genes that are most variable and shared across all samples. Post-integration, PCA was performed on the integrated data without additional scaling. Atlas-level UMAP embeddings were calculated using the first 20 principal components and Clustering was conducting as for individual datasets with a resolution parameter of 0.2.

Twelve different reference datasets (Liu_2022^7^; RuizMoreno_2022^25^; Nowakowski_2017^27^; Eze_2021^28^; Trevino_2021^29^; Aldinger_2021^30^; Bruggen_2022^31^; Braun_2023^32^; Velmeshev_2023^33^; Nano_2023^34^; Liu_2023^35^; Neftel_2019^36^; Table S3), encompassing various classifications and post-conception weeks (PCW), were utilized to correlate with the cellular states identified in this study. To achieve this, we used the top 50 markers previously identified for each cluster, together with the markers from the references, to calculate the signature scores across our dataset. Single-cell expression scores were computed following established methods^89,90^. For a given set of genes (Gj) representing an NMF program, metaprogram, or specific cell type, a score SCj(i) was calculated for each cell i, quantifying the relative expression of Gj. This was defined as the average relative expression (Er) of the genes in Gj compared to that of a control gene set (Gcont): SC(i) = average[Er(G, i)] − average[Er(Gcont, i)]. The control gene set consisted of the 100 genes with the most similar aggregate expression levels to each gene in Gj, ensuring a comparable distribution of expression levels while being 100-fold larger. Based on the scores computed for each signature across all references and cells, we then assessed the degree of correlation among them using a heatmap.

### Gene ontology analysis

Gene Ontology (GO) enrichment was carried out with g:Profiler2^91^ (v.0.2.2). For cancer or myeloid cell type, the top differentially expressed genes were calculated using Seurat’s ‘FindAllMarkers’ function as described above. Then, the ‘gost’ function was run using the following parameters: organism = “hsapiens”, correction_method = “gSCS”, evcodes=TRUE. Adjusted p-values were obtained using the g:SCS method, which controls for multiple testing by considering the structure of the ontology and the GeneRatio was calculated as intersection_size (genes that appear both in the input list and in the GO term) / effective_domain_size (all genes).

### Lineage scoring

Using the individual cell scores previously calculated for OPC-like, RG-like, iNPC/Neuronal-like, and GPC/AC-like cellular states, we performed the following analyses: cells were first partitioned into two groups — {OPC-like} / {RG-like} versus {iNPC/Neuronal} / {GPC/AC-like} — based on the sign of D, where D = max(SC_OPC_, SC_RG_) − max(SC_Neuronal_, SC_GPC/AC_). The value of D defines the y-axis coordinate for each cell. Subsequently, for {OPC-like} / {RG-like} cells (i.e., D > 0), the x-axis coordinate was defined as log_2_(|SC_OPC_ − SC_RG_| + 1). Conversely, for {iNPC/Neuronal-like} / {AC-like} cells (i.e., D < 0), the x-axis coordinate was defined as log_2_(|SC_Neuronal_ − SC_AC_|).

### Smart-seq2 processing

Raw sequencing reads from Smart-seq2 (SS2) experiments were aligned to the human reference genome (hg19) using HISAT2 (v.2.1.0), and gene-level quantification was performed with RSEM (v.1.3.0). For single-nucleus RNA-seq (snRNA-seq) data, the gene annotation files were modified to include intronic reads, allowing for quantification of both exonic and intronic signal. Following quantification, quality control was applied to retain high-quality SS2 cells. Cells from fresh samples were filtered based on the number of detected genes (nGene_cutoff > 1000) and mean expression of housekeeping genes (hk_mean_expr_cutoff > 2). For frozen samples, cells with more than 1,000 detected genes and an alignment rate greater than 30% (align_rate_cutoff > 30%) were kept.

To distinguish malignant from non-malignant cells, we applied inferCNV (v.1.20.0). Cells exhibiting tumor-like CNV patterns were classified as malignant and selected for downstream analyses, while diploid-like cells were excluded. Integration of the malignant SS2 cells was performed using the rliger R package (v.2.1.0), which implements integrative non-negative matrix factorization (iNMF). Gene expression profiles were normalized, scaled, and integrated using shared highly variable genes across samples. This approach identified five shared metagene factors that captured the major transcriptional programs within the tumor population. Annotation of SS2 tumor cells was achieved by scoring their expression profiles against cell states obtained from 10X data from this study. Each SS2 cell was assigned to the most similar 10X-derived cell state, enabling cross-platform consistency in downstream analyses.

### RNA velocity and pseudotime analysis

RNA velocity and pseudotime trajectories were inferred using the scVelo package^40^ (v.0.2.5) in a Python 3.9 environment. Spliced and unspliced transcript counts were quantified per sample using velocyto (v.0.17.17) to generate loom files. These files were subsequently merged and pre-processed using Scanpy (v.1.9.6), using previously defined cell state annotations. Transcriptional dynamics were modeled by applying scv.tl.recover_dynamics to estimate gene-specific kinetic parameters from the splicing kinetics of individual genes. RNA velocity vectors were then computed using the stochastic model (scv.tl.velocity) and embedded into a velocity graph via scv.tl.velocity_graph. The coherence of these vectors across neighboring cells was evaluated using scv.tl.velocity_confidence. To further quantify the magnitude of transcriptional change at the single-cell level, we calculated velocity length, defined as the Euclidean norm of the velocity vector in low-dimensional space. This metric served as a direct measure of the strength of dynamic transitions, highlighting cells with active progression through cellular trajectories. Velocity length was instrumental in distinguishing transitional states from more quiescent populations. Lastly, scv.tl.velocity_pseudotime was used to assign pseudotime values based on the directionality encoded in the RNA velocity vectors, enabling the reconstruction of temporal dynamics and lineage relationships in a fully unsupervised manner.

### Copy number variation analysis

CNV inference was performed using inferCNV (v1.22.0) with raw UMI count matrices extracted from the RNA assay. Cell-type annotations were derived from Seurat identities, and reference groups were specified as T cells (n = 300), B cells (n = 100), myeloid (n = 1000), endothelial cells (n = 600), pericytes (n = 400), and oligodendrocyte (OC) cells (n = 800). A fixed number of cells from each reference population was randomly sampled using a fixed random seed. Tumor cells were subsetted and grouped according to fusion status (NTRK1, NTRK2, NTRK3, MET, ROS1, and ALK). For tumor populations, all cells corresponding to the fusion of interest were retained. For ROS1 and ALK, tumor cells were further subset into two groups—(i) OPC-like and iNPC/neuronal-like and cycling progenitor cells, and (ii) radial glia-like and GPC/astrocyte-like cells—and analyzed separately to reduce computational burden and enable lineage-resolved CNV inference. Each fusion-specific dataset was analyzed independently. Gene coordinates were obtained from GENCODE v27 (hg38) and used to order genes along chromosomes. All chromosomes were retained for analysis. inferCNV was run using the following parameters unless otherwise stated: expression cutoff = 0.1, hierarchical clustering by cell groups enabled, denoising enabled, and hidden Markov model (HMM)-based CNV prediction enabled. Analyses were parallelized using eight computational threads. Final inferCNV outputs were used to generate heatmaps representing inferred copy number gains and losses across individual cells for each fusion group.

### SnATAC-seq data processing

Single-nucleus chromatin accessibility (scATAC-seq) data were generated using the 10x Genomics Chromium Single Cell Multiome ATAC + Gene Expression platform. Raw sequencing data were processed with Cell Ranger ARC (v.2.0) to generate fragment files and served as input for downstream analysis using ArchR^58^ (v.1.0.2). Each sample was analyzed independently. First an ArchRProject was created using the ‘createArrowFiles’ function and setting parameters to filter out low-quality cells based on a minimum transcription start site (TSS) enrichment score of 2 (filterTSS=2) and a minimum number of mapped ATAC-seq fragments required of 1’000 (filterFrags=1’000). Filtered fragments were then converted into a genome-wide ‘TileMatrix’ using 500-bp bins downstream analysis and a “GeneScoreMatrix” matrix was calculated. Next, potential doublets were identified and filtered out using the ‘addDoubletScores’ and ‘filterDoublets’ functions with default parameters. Dimensionality reduction and Clustering was performed using iterative Latent Semantic Indexing (LSI) on the “TileMatrix” with the following parameters (resolution=0.2, sampleCells=10’000, varFeatures=25’000, dimsToUse=1:30). Clustering was executed using the Seurat method from the LSI space with a resolution parameter of 0.8. UMAP embeddings were generated from the LSI space. For combined UMAP visualization across multiple samples, Harmony^92^ (v.1.2.0) was applied to correct for batch effects and was ran from the LSI space components.

We transferred cell type annotations from the matched snRNA-seq data. For downstream analysis, we only kept the cancer cells as annotated by snRNA-seq and merged all the data from the individually labeled samples together. To calculate average gene activity scores in the merged data and across cancer cell types, we use Seurat’s “FindAllMarkers” function on the “GeneScoreMatrix” with parameters set to a minimum percentage activity threshold (min.pct) of 0.5 and a log fold change threshold (logfc.threshold) of 0.5.

### Peak calling, and motif enrichment analysis, peak-to-gene linkage

Pseudo-bulk replicates were generated from the transferred cell-type annotation labels using ArchR’s internal algorithm. Peak calling was then performed using MACS2 (v.3.0.2) to generate a “PeakMatrix”. Motif annotations were then added to the peak set using the “addMotifAnnotations” function and using the CIS-BP database as reference^93^. We next identified marker peaks using ArchR’s “getMarkerFeatures” function with a Wilcoxon test. We then used ArchR’s “peakAnnoEnrichment” (setting peakAnnotation = “Motif”) to reveal which motifs were enriched among these marker peaks by comparing the observed motif occurrences to a background set. This yields a motif enrichment score for each cell type across all samples. In addition, we used also “peakAnnoEnrichment” (setting peakAnnotation = “ATAC”) to project our identified peaks onto a reference bulk ATAC-seq dataset.

We used ArchR’s ‘addPeak2GeneLinks’ function to infer regulatory links between peaks and genes. Briefly, ArchR first defines accessible peaks (via pseudo-bulk peak calling using MACS2). Next, for each peak in the genome, ArchR computes correlation between that peak’s accessibility and the gene’s accessibility (inferred from “GeneScoreMatrix”) across pseudo-bulk replicates. The strength of correlation is then used as evidence for a putative regulatory link. The result is a “Peak2Gene” linkage score that identifies potential cis-regulatory interactions driving gene expression.

### Epigenetic plasticity score

To assess regulatory plasticity, we calculated a per-cell entropy-based score using ATAC-derived gene activity scores across the RNA-defined cancer subpopulations. Top gene activity scores were identified from differential accessibility analysis from the “GeneScoreMatrix” using Seurat’s “FindAllMarkers” as described above. Cells were then scored for each cancer cell type using Seurat’s AddModuleScore. The resulting values were transformed into probability distributions using a softmax function (temperature = 0.5). Shannon entropy was then computed per cell and normalized by the theoretical maximum entropy (log K, with K = 5) to yield a plasticity score ranging from 0 (regulatory commitment) to 1 (maximal epigenetic plasticity).

### Gene regulatory network inference

Gene regulatory networks (GRNs) were modeled per sample using Pando^55^ (v.1.0.0), in a merged single-cell dataset, which merged either RG/GPC/AC-like, OPC-like, or iNPC/Neuronal-like cell types from three samples (PT0101, PT0201, PT0301). Candidate regulatory regions were defined by the peaks identified using MACS2 in the ATAC-seq data using the “initiate_grn” function. These peaks were then combined with SCT-normalized expression values into a SeuratPlus object per cell type. Transcription factor binding motifs were scanned within these candidate regions using the ‘find_motifs’ function. GRN inference was performed using the ‘infer_grn’ function, focusing on the variable genes identified from SCT-normalized RNA-seq values. The peak-to-gene linkage was determined using the “GREAT” method. Inferred GRN modules were finally ranked based on the number of target genes in their modules and visualized either as a ranked list or graph.

### MERFISH image preprocessing and processing

MERFISH data were acquired using the MERSCOPE platform from Vizgen. Raw data included high-resolution images and corresponding spatial transcript counts. The analysis was conducted using the SOPA (Spatial Omics Processing and Analysis) pipeline^94^ (v.1.1.1). Briefly, due to the large size of the imaging data, images were patched into 6000x6000px tiles with an overlap of 150px. Cell segmentation was then performed using two algorithms: Cellpose^72^ and Baysor^73^ . Cellpose was used to segment cells based on the DAPI staining, while Baysor was used to segment cells based on spatial transcriptomic data, while using Cellpose prior segmentation information. Post-segmentation, transcript counts were aggregated within cell segmentation boundaries. Transcript counts and x,y coordinates were then loaded into a Seurat object as a ‘SOPA’ assay for further processing. Briefly, cells with n<25 genes or n<50 transcripts identified were removed. SCT normalization on all transcripts, followed by PCA and UMAP visualization were performed.

### Xenium in situ imaging

Image acquisition and sample handling was automated within Xenium Analyzer for two slides per run. Fifteen rounds of fluorescence probe hybridization, imaging and probe removal occurred within the Analyzer. A fast area scan camera with ∼200 nm per pixel resolution was used for image acquisition, and z stacks were taken with 0.75 μm step size across tissue thickness. All z-stack of images were then processed and stitched, using DAPI image as reference. Each fluorescently labelled oligonucleotide bound to amplified barcodes was detected and registered in each cycle. Unique optical signature from fluorescence intensity over the 15 rounds was used to identify a target gene. For cell segmentation, neural network was used to detect each nucleus from DAPI images.

### Preprocessing 10x Xenium Samples

For processing 10x Xenium data, standard preprocessing steps from Seurat (v5.3.0) were used. First, the raw output from Xenium Analyzer was loaded using LoadXenium() function. Then, the data was normalized using SCTransform (v0.4.2). This was followed by dimension reduction using Principal Component Analysis, projection into 2 dimensions using UMAP, and construction of neighborhood graph followed by unsupervised clustering of cells.

### Cell state/type annotation in spatial samples

A similar strategy was used to annotate both MERFISH and Xenium samples. The cell state/type annotation in spatial samples was done in a 2-step process. In the first step, we annotate the normal cell types in the sample. We use the 10x single-cell data from Project CARE (The multilayered transcriptional architecture of glioblastoma ecosystems, Nomura et al. (2025)) for this purpose. First, for the MERFISH/Xenium samples, the single-cell data is subsetted to just the genes in the MERFISH/Xenium panel to yield two separate objects. Then, both the resultant single cell objects are freshly processed using standard pipeline from Seurat (v5.3.0), which includes: i) normalization using SCTransform (v0.4.2), ii) dimension reduction using Principal Component Analysis, iii) batch correction using harmony (v1.2.3) iv) projection into 2 dimensions using UMAP.

Then, a label transfer is performed from the appropriate single cell object onto the data for each spatial sample (MERFISH/Xenium). For the label transfer, we use the anchor-based label transfer approach from seurat, which involves using the functions FindTransferAnchors() and TransferData(). This leads to the normal celltypes being annotated in all the spatial samples, and remaining cells being called “Malignant” at this stage.

In the second step, we perform a second label transfer onto the cells annotated as “Malignant” in the spatial samples till this point. We take the 10x single cell data (with finer malignant metaprogram annotation) we generated in this study, and similar to the procedure for normal cells’ annotation, subset to just the genes in MERFISH/Xenium panels to obtain two single cell objects. Then process both those objects exactly like the single cell references with normal cells (above) to make them ready for label transfer. Finally, onto the “Malignant” only subset of spatial samples, we perform label transfer (anchor-based approach from Seurat) from the appropriate single cell reference with finer malignant annotations.

### Neighborhood analysis

We used the python package squidpy (v1.6.5) to quantify the significance of a celltype pair neighbors, whose centroids are within a distance of 20um from each other. Then, we used the function squidpy.gr.nhood_enrichment() to compute the neighborhood enrichment of each celltype pair using a permutation test with 1000 permutations. Finally, to identify recurring patterns across all the samples, we averaged the neighborhood enrichment z-score values across all the samples.

### Niche and Metaniche analysis

We used BuildNicheAssay function from Seurat to identify niches with k=6 in each sample. This function computes the number of cells of different types in the neighborhood (defined by 20 nearest neighbors) of each cell and uses k-means clustering to group the cells into k niches. To identify recurring niches across samples, we computed celltype proportions for each niche in each sample, and computed pairwise pearson correlation between all of them. Then, we performed hierarchical clustering to cluster all the niches across all samples and used the cutree function from stats (v4.4.2) package in R to obtain n=6 groups of niches, which we call “metaniches”.

### Reporting summary

Further information on research design is available in the Nature Portfolio Reporting Summary linked to this article.

### External data collection

Bulk RNAseq and methylation data (FASTQ, BAM, IDAT files) was requested for patients PT0101, PT0103, PT0104, PT0105, PT0106, and PT0107 from the Children’s Brain Tumor Network. MGF generated the scRNA-seq data (Smart-seq2). ADM and COAB performed the snRNA-seq bioinformatic analyses, and ADM performed the snATAC-seq and MERFISH analyses with contributions from CM, FZ, TR, and RR. Primary tissue resources and additional data interpretation were provided by MZ, LN, UT, CH, CJ and JG. AGS and MGF supervised all aspects of the study.

cooccuring within each other’s vicinity in each sample. We used the function squidpy.gr.spatial_neighbors() with coord_type = ‘generic’ and radius = 20, to construct a neighborhood graph according to spatial coordinates of cells, defining any two cells as

## Data availability

SnRNA-seq, snATAC-seq, and scRNA-seq (Smart-seq2), and bulk RNA-seq data of primary patient RTK-fused gliomas have been submitted to GEO. Previously published data used in this study are available under accession codes: GSE184357^7^, GSE162989^11^, GSE211376^25^, phs000989.v3^27^, SCR_015820^28^, GSE162170^29^, phs001908.v2.p1^30^, EGAS00001006136^31^, EGAS00001004107^32^, nemo:dat-3ah9h9x^33^, GSE242779^34^, BioProject:PRJNA798712^35^, GSE131928^36^.

## Acknowledgments

This work has been supported by the Swiss National Science Foundation (#PCEFP3_194620), Swiss Bridge Award, Promedica Stiftung awarded to AGS, as well as Universität Zürich Postdoc Grant awarded to ADM. This work was also supported the NIH Director’s New Innovator Award (DP2NS127705), the Distinguished Scientist Award from the Sontag Foundation and Caroline Mortimer Fund awarded to M.G.F. and Name]. C.L.C is a Damon Runyon-St. Jude Pediatric Cancer Research Fellow supported by the Damon Runyon Cancer Research Foundation and St. Jude Children’s Research Hospital (SJP-#01-24). R.R.M. is supported by The Brain Tumour Charity Future Leaders Fellowship (FL_2025_/1_11049), and the Filling the Gap grant from the UZH. The authors would like to acknowledge the staff at the Princess Margaret Genomics Centre at UHN Toronto (Johana Regala, Samantha Lee, Julissa Tsao, Wafa Al-Ameri, and Dr. Troy Ketela) for their unwavering support in generating high-quality single-cell and transcriptomic data. The authors would also like to thank Dr. Antonela Petrović for constructive feedback on the results and manuscript.

## Author contributions

ADM, COAB, MGF, AGS conceived the study and designed the experiments. ADM, COAB, CLC, CM RM, VK, SK, AP, FMGC, MGF, AGS interpreted results. ADM, COAB, CLC, MGF, and AGS wrote the first draft manuscript with the input of all co-authors. ADM and AGS generated the snRNA/ATAC-seq (10X Multiomics), bulk RNA-seq, methylation, and spatial transcriptomic data (MERFISH). COAB and MGF generated the scRNA-seq data (Smart-seq2). ADM, COAB and CM and performed the snRNA-seq bioinformatic analyses with support from VK, and ADM performed the snATAC-seq analyses with contributions from CM, FZ, TR, and RR. Xenium and MERFISH analysis were performed by SK and ADM, with contributions from COAB, CLC and CM. Primary tissue resources and additional data interpretation were provided by MZ, LN, UT, CH, CJ and JG. AGS and MGF supervised all aspects of the study.

---

**Figure S1.**
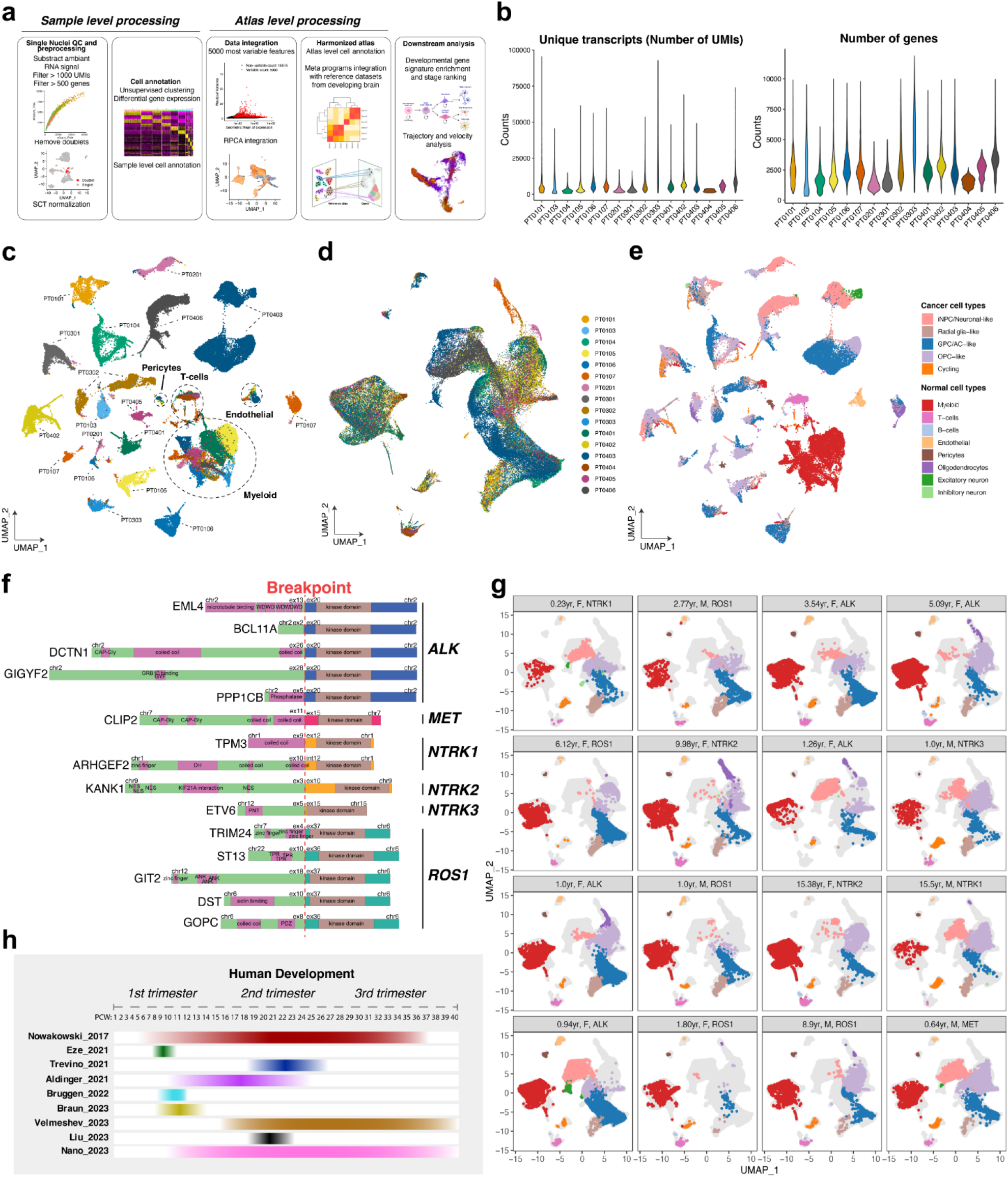
Single-nucleus RNA-seq analysis pipeline. (a) Data processing workflow. (b) Distribution of the number of unique transcripts (UMIs) and number of genes identified in individual nuclei per patient sample. (c) UMAP of unintegrated data (no batch correction) colored by patient. (d) UMAP of batch corrected data colored by patient. (e) UMAP of unintegrated data colored by cell type. (f) Main patient RTK gene fusion and breakpoints. (g) Integrated UMAP embeddings of single-nucleus transcriptome from individual patients (10X Multiome, n=16) and color-coded by cell type. (h) Developmental datasets used for data integration (see also Fig. S3) with the corresponding developmental windows

**Figure S2.**
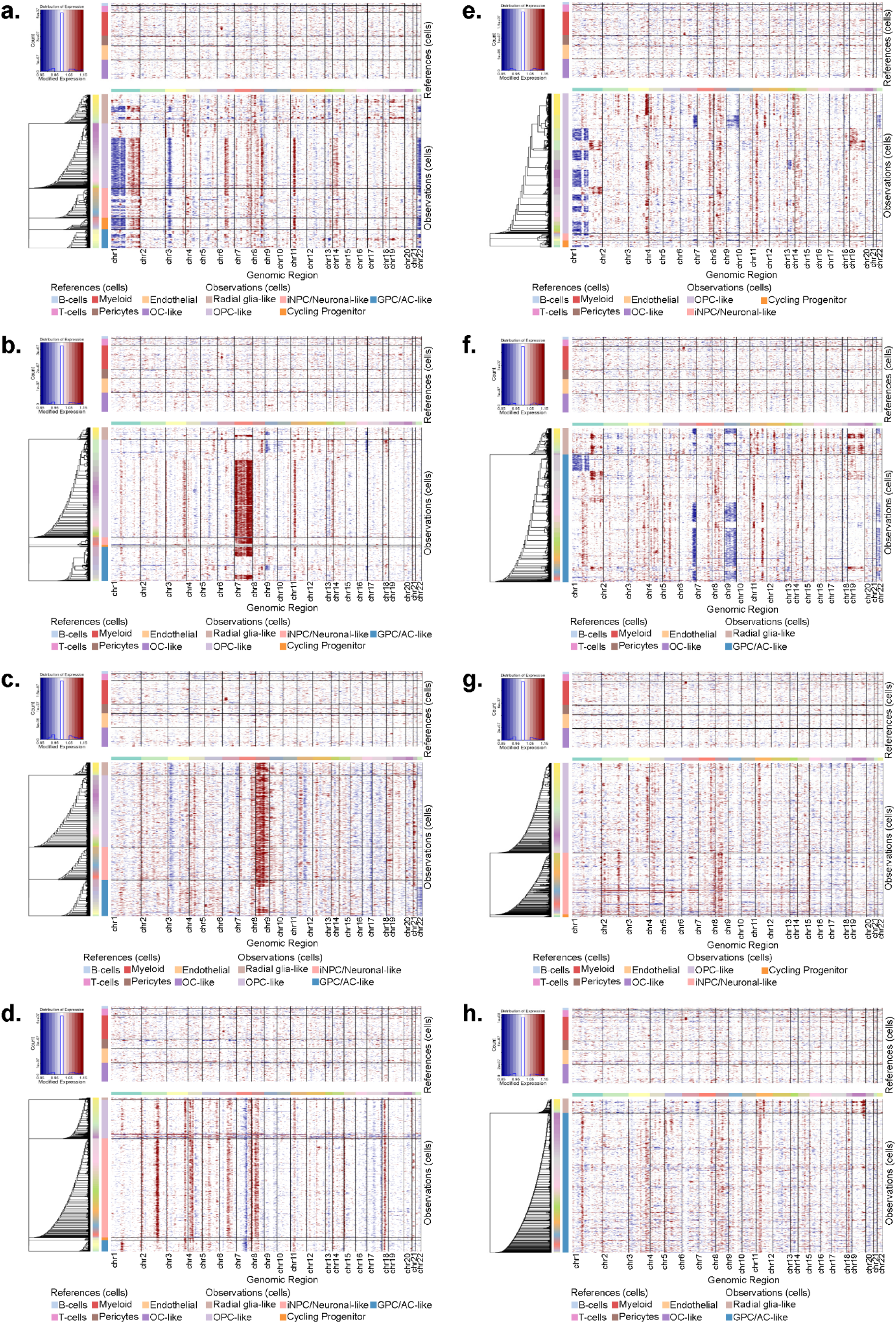
Chromosomal copy number variation inferred from single-cell RNAseq. For all analyses, inferCNV reference profiles were constructed using T cells, B cells, myeloid, endothelial cells, pericytes, and oligodendrocyte-like (OC-like) cells. Heatmaps indicate inferred copy number gains and losses across individual cells. (a) NTRK1 samples. (b) NTRK2 samples. (c) NTRK3 samples. (d) MET samples. (e) ROS1 samples restricted to OPC-like, iNPC/Neuronal-like cells and Cycling Progenitor cells. (f) ROS1 samples restricted to radial glia-like and GPC/astrocyte-like cells. (g) ALK samples restricted to OPC-like, iNPC/Neuronal-like cells and Cycling Progenitor cells. (h) ALK samples restricted to radial glia-like and GPC/astrocyte-like cells.

**Figure S3.**
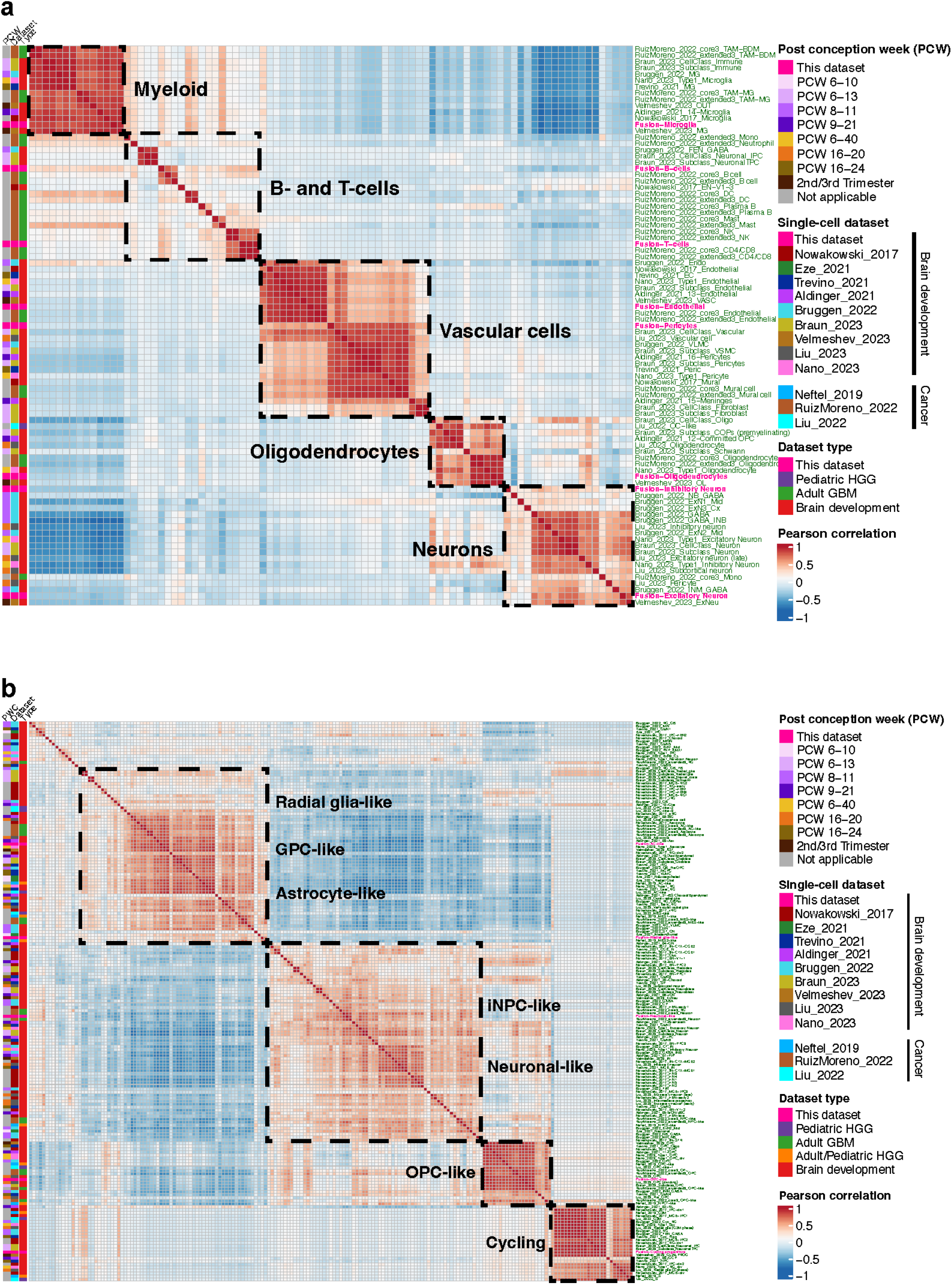
Single-cell data integration. Heatmap showing Pearson correlation scores between normal (a) and cancer (b) cell gene signatures from our dataset (pink labels) and published single-cell signatures of the developing brain and brain cancer (green labels).

**Figure S4.**
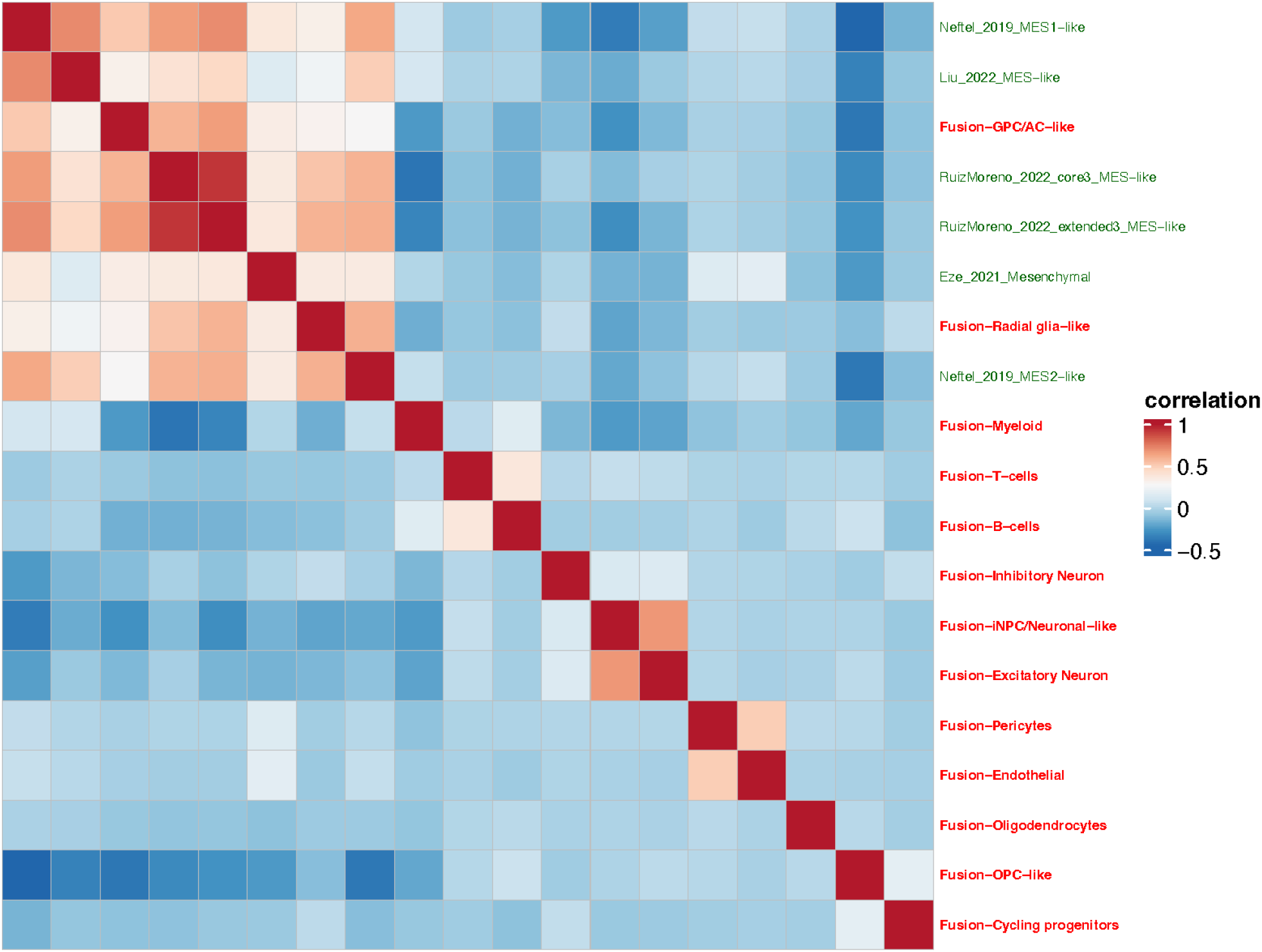
MES-like signature correlation. Heatmap showing Pearson correlation scores between all cells from our dataset (red labels) and MES-like populations from publicly available datasets (green labels).

**Figure S5.**
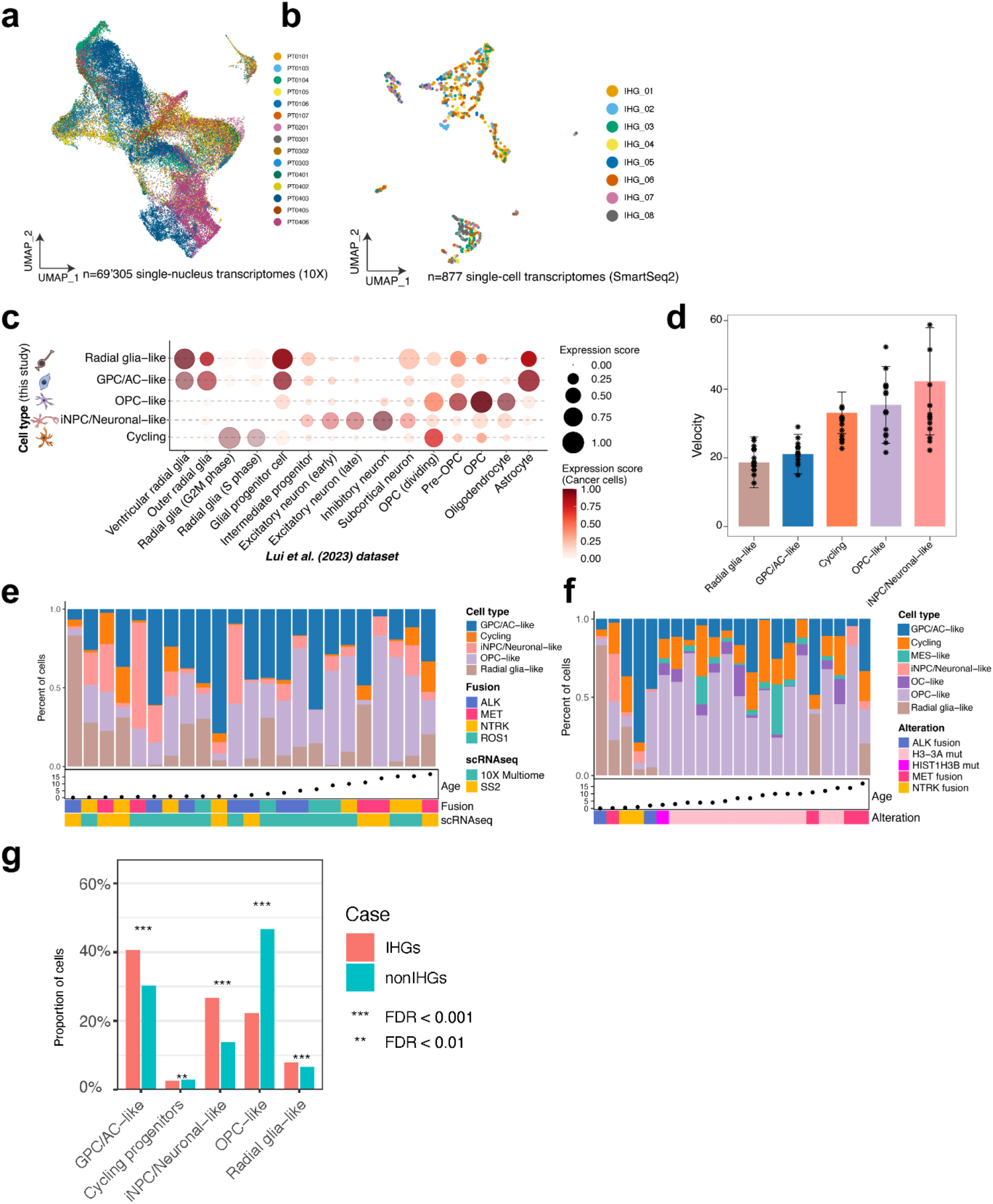
SmartSeq2 atlas of IHGs and cancer cell population heterogeneity. (a) UMAP of 69’305 single-nucleus transcriptomes colored by patient. (b) NMF-integrated UMAP of 877 cancer single-cells transcriptomes profiled by SmartSeq2 colored by patient. (c) Projection of cancer metaprograms onto Liu et al. (2023) developing brain single-cell atlas. Color scale: scores of normal cell signatures in cancer cells. Dot sizes: scores of cancer cell signatures in normal cells. (d) Average velocity value per sample and across cancer metaprograms. (e) Proportion of cancer cell types across the cohort (10X Multiome and SmartSeq2) and ordered by age. (f) Bar plot representing the proportion of tumor cell populations in SmartSeq2 datasets of RTK-driven gliomas (Fig. 2E) and H3K27M diffuse midline gliomas (Liu et al. (2024)). (g) Ratios of cell counts between the RG-like and GPC/AC-like populations and iNPC/Neuronal-like populations for n=8 infant patients ordered by age. Black bars for deceased patients. (h) Distinct cellular compositions of IHGs and non-IHG gliomas. Proportions of malignant cell populations identified by SmartSeq2 profiling in IHGs and non-IHG gliomas. Bars represent the percentage of cells belonging to each population within each tumor group. Statistical significance of differences in cell-state proportions between groups was assessed using differential proportion analysis with false discovery rate (FDR) correction; ***FDR < 0.001 and **FDR < 0.01.

**Figure S6.**
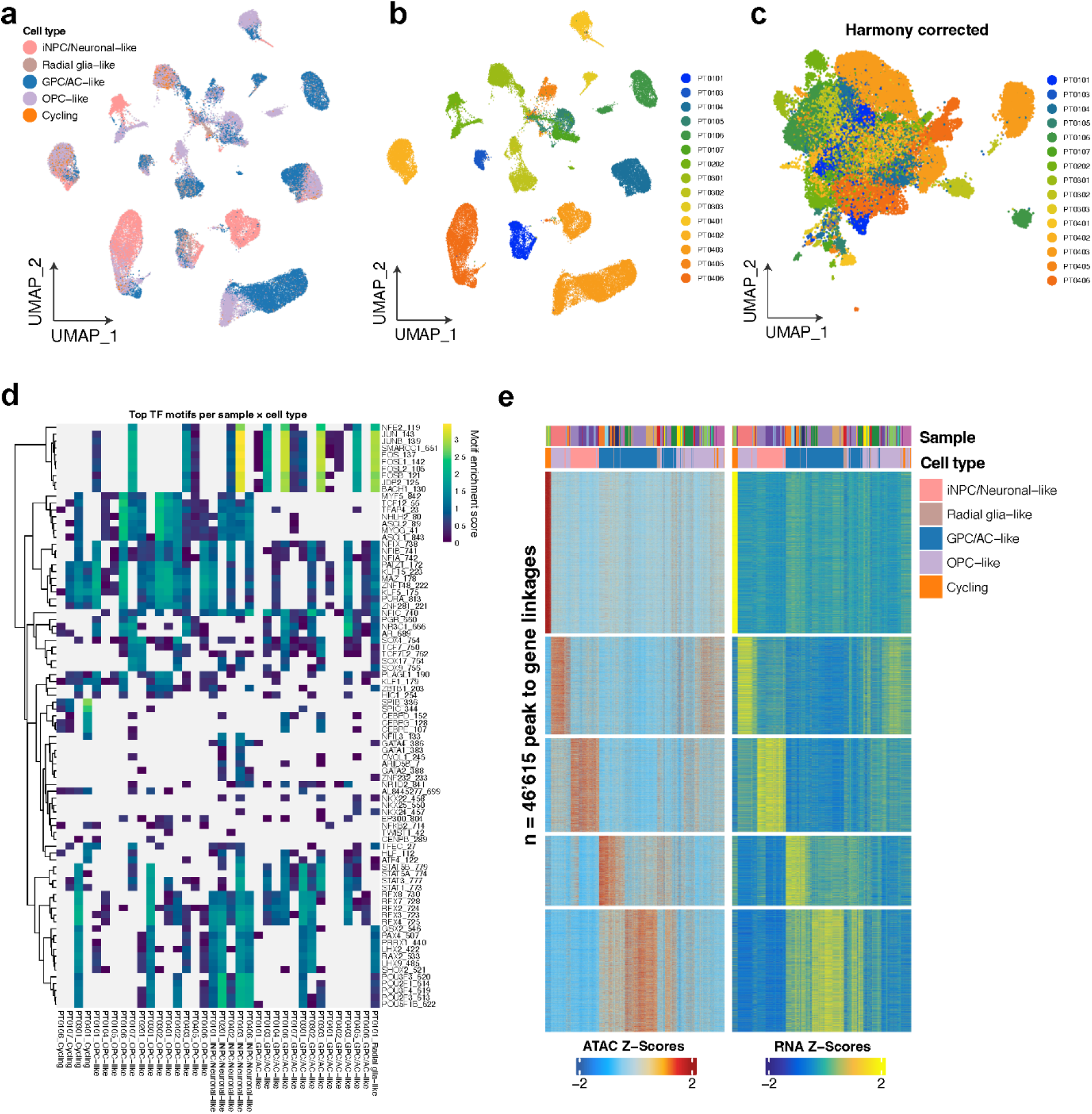
snATAC-seq atlas and chromatin accessibility analysis. (a) Unintegrated snATAC-seq atlas of n=15 patients colored by cancer cell type. (b) Unintegrated snATAC-seq atlas of n=15 patients colored by patient. (c) Harmony-corrected (integrated) snATAC-seq atlas of n=15 colored by patient. (d) Motif enrichment analysis on differential peaks across all patients and tumor cell type. Grey cells indicate motifs that were not detected. Only populations with > 30 cells were considered for this analysis. (e) Heatmap of peak-to-gene links displaying normalized gene expression and chromatin accessibility. A total of n=46’615 significantly linked peaks were identified. Paris (row) clustered using k-means (k=5). Legend represents the sample and annotated cell type.

**Figure S7.**
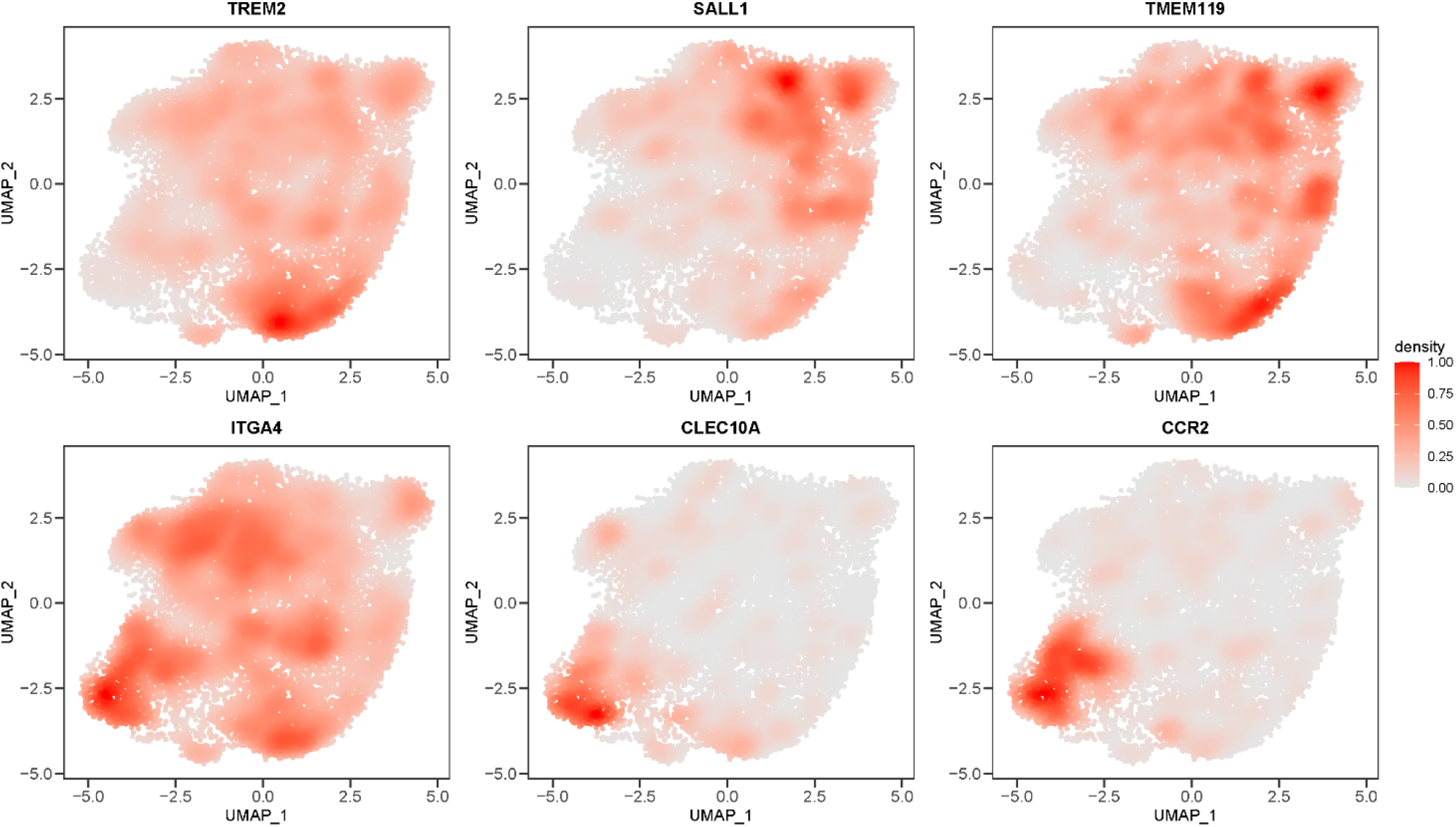
Tumor-associated macrophages. UMAP density plots of macroglia- and macrophage-associated transcripts.

**Figure S8.**
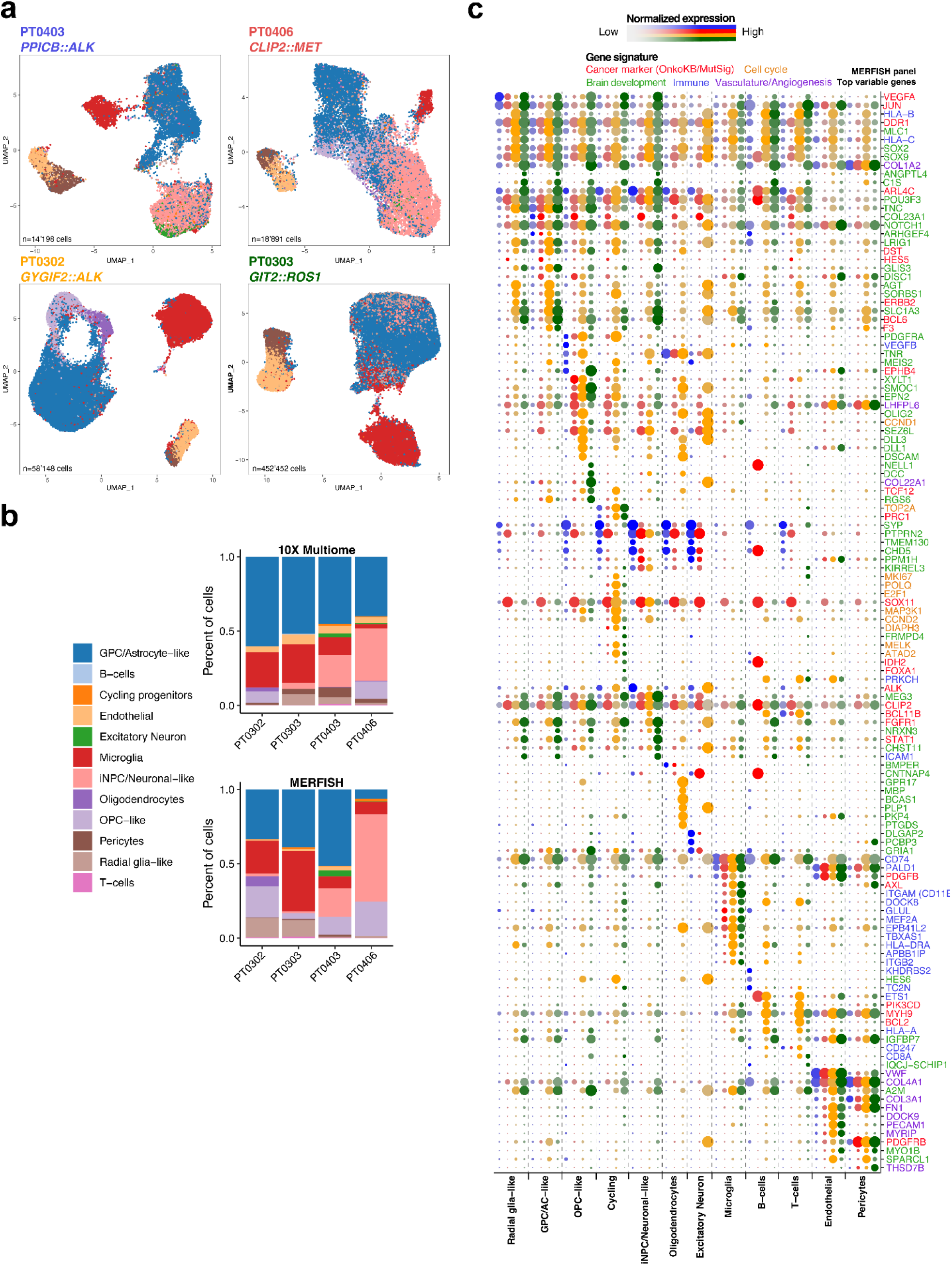
MERFISH patient atlases, cell type composition, and top variable genes. (a) UMAP of single cells resolved by MERFISH after QC filtering (n=500 genes) and colored cell type. (b) Proportions of cell types across patient samples, profiled by either 10X Multiome or MERFISH (inferred cell types for MERFISH only). (c) Dot plot of gene expression for top variable markers colored by patient (Fig. S6A) that were identified as differentially expressed across the tumor cell populations. Genes colored by marker type including cancer, brain development, cell cycle, immune, and vasculature, angiogenesis. Dot size indicate the percentage of cells expressing each per sample, while color intensity represents normalized expression level.

